# Spatially confined sub-tumor microenvironments orchestrate pancreatic cancer pathobiology

**DOI:** 10.1101/2021.02.18.431890

**Authors:** Barbara T Grünwald, Antoine Devisme, Geoffroy Andrieux, Foram Vyas, Kazeera Aliar, Curtis W McCloskey, Andrew Macklin, Gun Ho Jang, Robert Denroche, Joan Miguel Romero, Prashant Bavi, Peter Bronsert, Faiyaz Notta, Grainne O’Kane, Julie Wilson, Jennifer Knox, Laura Tamblyn, Nikolina Radulovich, Sandra E Fischer, Melanie Boerries, Steven Gallinger, Thomas Kislinger, Rama Khokha

## Abstract

Pancreatic ductal adenocarcinoma (PDAC) remains resistant to most treatments and demonstrates a complex pathobiology. Here, we deconvolute regional heterogeneity in the human PDAC tumor microenvironment (TME), a long-standing obstacle, to define precise stromal contributions to PDAC progression. Large scale integration of histology-guided multiOMICs with clinical data sets and functional *in vitro* models uncovers two microenvironmental programs in PDAC that were anchored in fibroblast differentiation states. These sub-tumor microenvironments (subTMEs) co-occurred intratumorally and were spatially confined, producing patient-specific cellular and molecular heterogeneity associated with shortened patient survival. Each subTME was uniquely structured to support discrete aspects of tumor biology: reactive regions rich in activated fibroblast communities were immune-hot and promoted aggressive tumor progression while deserted regions enriched in extracellular matrix supported tumor differentiation yet were markedly chemoprotective. In conclusion, PDAC regional heterogeneity derives from biologically distinct reactive and protective TME elements with a defined, active role in PDAC progression.

**Graphical Abstract & Key findings:** 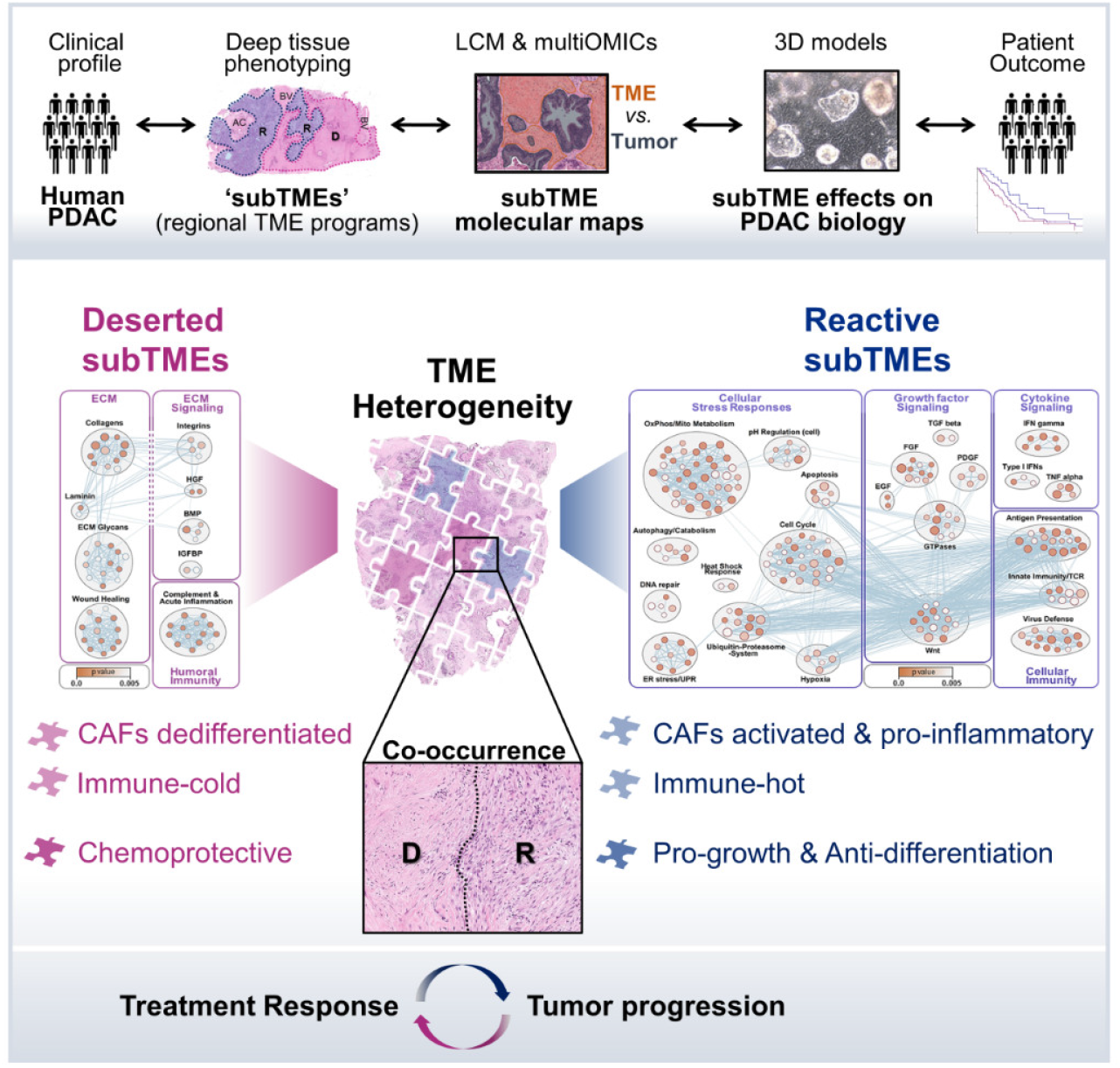

- PDAC regional heterogeneity originates in sub-tumor microenvironments (subTMEs)
- SubTMEs exhibit distinct immune phenotypes and CAF differentiation states
- Different subTMEs are either tumor-promoting or chemoprotective
- Intratumoral subTME co-occurrence links stromal heterogeneity to patient outcome

## Introduction

The dismal prognosis of pancreatic ductal adenocarcinoma (PDAC) and a the recurrent trend of clinical trial failure (Rahib et al., 2016) urge a better understanding of this complex malignancy. Among its most recognised features is the abundant stromal reaction, an ecosystem of immune cells, endothelial cells and cancer-associated fibroblasts (CAFs) that provides specialized niches for cancer cells and modulates tumor growth and invasive behaviour (Ho et al., 2020). Still, discrepancies remain on how to annotate relevant stromal elements and their respective pro- or anti-tumor functions. For example, recent work revealed clinically relevant subtypes of PDAC stroma (Maurer et al., 2019; Moffitt et al., 2015; Puleo et al., 2018) and showed that ECM-rich/desmoplastic transcriptional signatures are associated with a worse prognosis. Yet, tissue-based studies suggest that collagen- and ECM-rich stroma yields a better prognosis (Bolm et al., 2017; Erkan et al., 2008). Similarly, despite the wealth of functional studies showing that CAFs can promote PDAC progression and chemoresistance (Ligorio et al., 2019; Ogawa et al., 2021), higher stromal content can yield a better prognosis (Knudsen et al., 2017; Torphy et al., 2018) and the depletion of CAFs can result in more aggressive tumors in mice (Özdemir et al., 2015; Rhim et al., 2014). Thus, interactions between PDAC and stromal cells are likely not uniformly tumor promoting or inhibitory and the underlying intricate nuances have remained poorly understood.

Recent landmark studies in the field focused on single cell types revealed the remarkable cellular complexity of the PDAC tumor microenvironment (TME), comprising numerous stromal cell subpopulations, especially in the CAF compartment (Biffi et al., 2019; Dominguez et al., 2020; Elyada et al., 2019; Öhlund et al., 2017). In addition, we are learning that PDAC displays a unique immune milieu with heterogenous immunosuppressive populations suspected to hamper current immunotherapeutic strategies (Fan et al., 2020). These complex cellular communities and their intricate interactions in the PDAC TME encompass a myriad of possibilities for how they could be sustaining growth and progression of the malignant epithelium. A major obstacle for assigning clear functions to the TME in PDAC lies in its inter- and intra-tumoral heterogeneity (Iacobuzio-Donahue et al., 2020; Vickman et al., 2020), which has implications for bulk analyses and sampling but also suggests the presence of distinct and possibly independent regional programs in the PDAC TME. Given that histology is reflective of how cell communities in a tissue self-organize to establish their function, we hypothesized that recurrent phenotypes within the heterogenous TME could signify stromal ‘functional units’ relevant for PDAC pathobiology.

## Results

### Widespread subTMEs in human PDAC are associated with poor survival

We aimed to find regional histological patterns in the PDAC TME that recurred across patients, stages, and sites. Regional TME heterogeneity was largely due to variations in cellular and acellular stromal components, the maturation and extent of ECM, and differences in stromal cell morphology (**Figure 1A**). Anchored on these histological features, we reviewed multiple whole sections per PDAC resection (n = 143) and core needle biopsies from advanced PDAC cases (n = 210; **Figure 1B, Table S1**). This comprehensive review identified three recurrent TME phenotypes in PDAC (**Figure 1C**): 1. deserted regions with thin, spindle-shaped fibroblasts embedded in loose and fully matured collagen fibers, often with keloid and myxoid features; 2. reactive regions containing fibroblasts with plump morphology and enlarged nuclei, rich in inflammatory infiltrate and acellular components; 3. in addition to these two ‘extremes’, we recorded regions with intermediate levels of these features. These three TME phenotypes (deserted, intermediate, reactive) were also observed at other common metastatic sites (**Figure 1D**). Assessment of the dominant TME phenotype (>50% area contribution; **Figure S1A**) showed that these three TME phenotypes occurred widely tumors across PDAC stages (**Figure 1E**) and cohorts (TCGA PAAD cohort (TCGA Research Network, 2017); **Figure 1F**).

**Figure 1.**
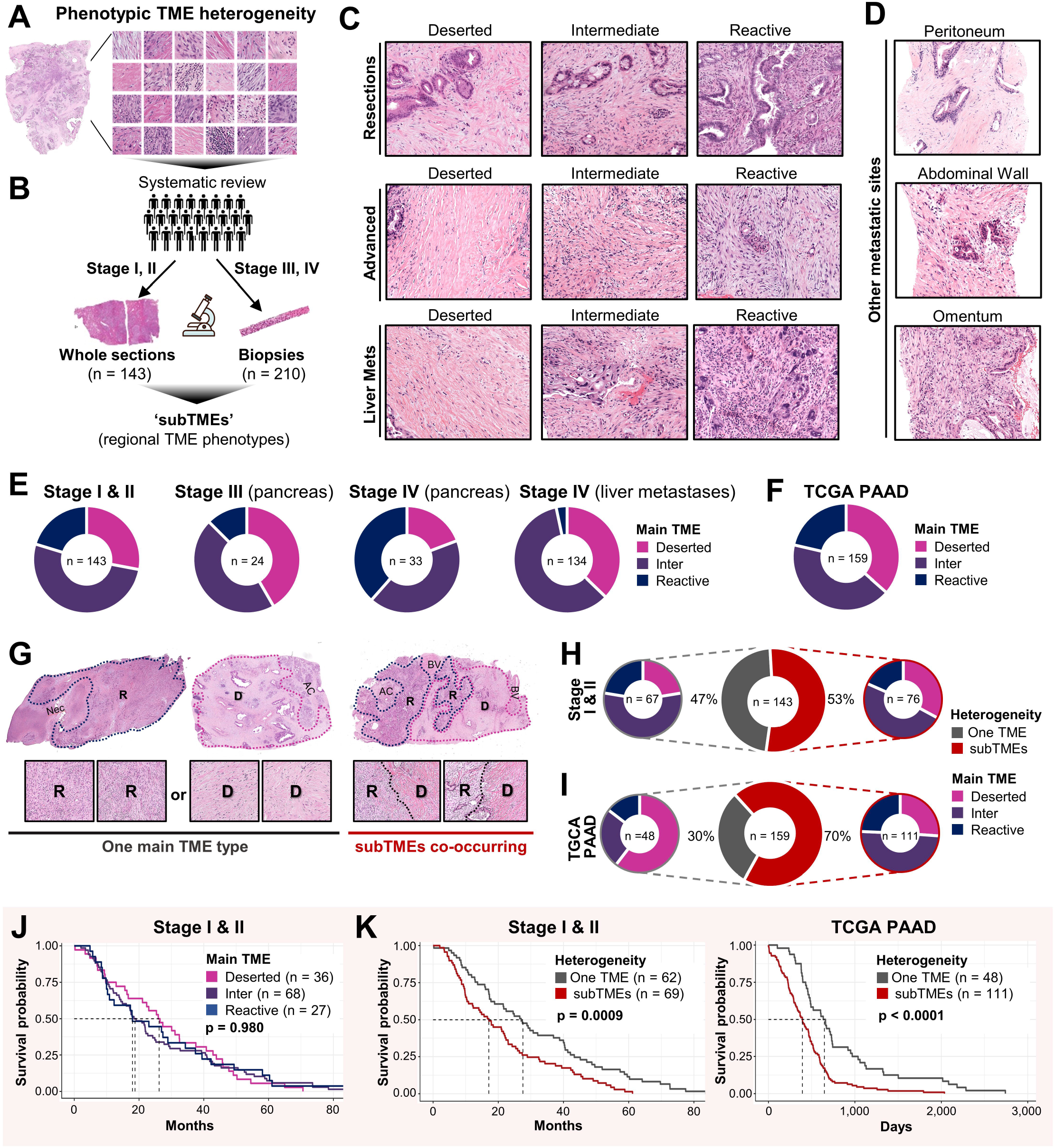
Widespread subTMEs in human PDAC are associated with poor survival. See also Figure S1 and Table S1. **A)** Schematic: Regional phenotypic heterogeneity in the human PDAC TME. Shown are multiple regions (*right*) from one PDAC tumor whole section (*left*). **B)** Review strategy for identification of regional TME phenotypes (= sub-tumor microenvironments (subTMEs)) in human PDAC across disease stages and sites. **C-D)** HE images of the deserted, intermediate, and reactive subTME phenotypes **C)** in resectable primary tumors (*top*), advanced primary tumors (*middle*), liver metastases (*bottom*), and **D)** other common metastatic sites. **E, F)** Frequencies of main TME phenotype (= subTME with >50% area contribution) in **E)** Toronto in-house cohorts and **F)** the TCGA PAAD cohort. **G)** Representative images of 2 cases with one reactive or deserted main TME phenotype (*left*) *vs*. a case with co-occurring reactive and deserted subTMEs (*right*). D, deserted subTME; R, reactive subTME; Nec, necrosis; BV, blood vessel; AC, acini. **H-I)** Nested donut plots showing frequencies of subTME co-occurrence (*centre donut*). Nested donut plots show main TME frequencies for cases with one main TME (grey subgroup, *nested left*) *vs*. co-occurring subTMEs (red subgroup, *nested right*) in **H)** the resection cohort and **I)** the TCGA PAAD cohort. **J-K)** Kaplan-Meier analysis of overall survival. Log rank tests. **J)** Resection cohort; strata: main TME phenotype. **K)** Resection cohort (left) and TCGA PAAD cohort (*right*); strata: single dominant TME phenotype (‘One TME’) *vs*. subTME co-occurrence (‘subTMEs’).

Noticeably, the different phenotypes could co-occur within the same tumor in a spatially defined manner (**Figure 1G**), which led us to name them ‘sub-tumor microenvironments’ (subTMEs). This phenomenon was frequent: co-occurrence of subTME phenotypes (**Figure S1B**) was observed in 76/143 (53%) of primary tumors in our resection cohort (**Figure 1H**) and 111/159 (70%) in the TCGA PAAD cohort (**Figure 1I**), irrespective of the dominant subTME in the tumor (**Figure 1H-I**, nested plots). Importantly, patient overall survival was not linked to the dominant TME phenotype (**Figure 1J, Figure S1C**) but to the intratumoral co-occurrence of subTMEs (**Figure 1K**), i.e. a phenotypically heterogenous TME. In sum, we categorized the human PDAC TME into distinct regional phenotypes termed ‘subTMEs’. These occurred across disease stages and sites and had a profound combinatory effect on patient survival.

### Molecular profiling of subTMEs and patient-matched malignant epithelium in PDAC

To study the composition and function of subTMEs, we assembled a workflow for large-scale patient-matched integration of histopathology, clinical datasets, multidimensional analytical techniques, and subTME-specific model systems. Our first step was to construct subTME-specific multiOMIC profiles from 32 of the 143 patients in our Stage I&II cohort (**Figure 2A**) with typical tumor morphology, sufficient material, sufficient stromal content (> 50%), matched high-quality whole genome sequencing (WGS), and no homologous recombination-repair deficiency (Golan et al., 2021) or other rare genomic events. We used laser capture microdissection (LCM) to separately extract different subTMEs and the tumor cells residing within (**Figure S2A**). Samples were split for parallel profiling of the proteome via shotgun proteomics and the transcriptome by RNAseq. This was successful for 29/32 patients with 3/32 patient yielding only enough material for proteome analysis. MultiOMIC profiling then identified 5,708 unique protein groups (**Figure 2B**) and 17,945 transcripts (**Figure 2C**). Enrichment of TME and tumor epithelial fractions by LCM was highly effective: tumor *vs*. TME samples separated widely in principal component analysis (PCA; **Figure S2B**) and showed distinct expression profiles with 5,070 differentially expressed genes (DEGs) at the proteomic level and 13,071 DEGs at the transcriptomic level (**Figure S2C**), including known TME markers ACTA2 (proteome) and PDGFRA (transcriptome) in the top hits. We furthermore observed a near-complete enrichment of recent thoroughly validated LCM-based stromal and epithelial RNA gene signatures (Maurer et al., 2019) by GSEA (**Figure S2D**). In sum, we built detailed multiOMICs profiles of different PDAC subTMEs and the inhabiting malignant epithelium.

**Figure 2.**
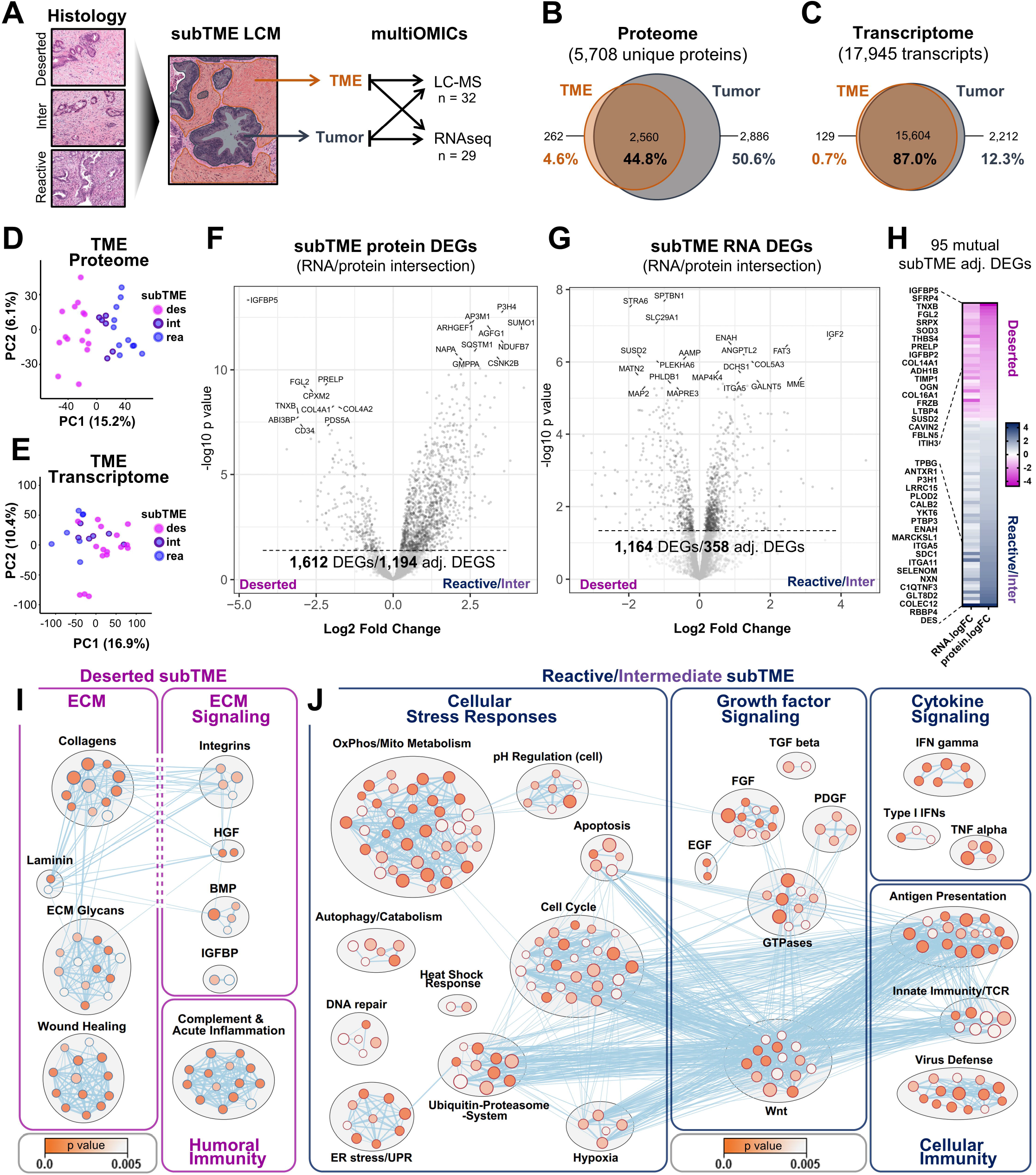
SubTME molecular landscapes differ in ECM, CAF activation, and immune features. See also **Figure S2** and **Tables S2-S4** **A)** Schematic: Histology-guided laser capture microdissection of subTMEs and patient-matched tumor epithelium from human PDAC tissue for parallel proteomic and transcriptomic profiling. **B-C)** Venn diagrams of genes detected in the TME and in the tumor fraction of human PDAC subTME-guided **B)** proteomes and **C)** transcriptomes. **D-E)** Principal Component Analysis (PCA) of TME fractions from **B-C)**, for **D)** proteomes and **E)** transcriptomes. Colours indicate subTMEs. **F-G)** Volcano plots: differentially expressed genes (DEGs) in deserted *vs*. reactive/intermediate subTMEs at **F)** protein level and **G)** RNA level following proteome-transcriptome integration. **H)** Heatmap depicting fold changes (fc) for adjusted DEGs with mutual regulation on RNA and protein level in deserted *vs*. reactive/intermediate subTMEs. **I-J)** Enrichment map summarizing gene sets up-regulated in **I)** deserted subTMEs (see also **Table S3**) and **J)** reactive subTMEs (see also **Table S4**).

### SubTME molecular landscapes differ in ECM, CAF activation, and immune features

We proceeded to interrogate the TME fractions of the multiOMIC datasets. Remarkably, proteomes separated by subTME on the first principal component (**Figure 2D**) and exhibited vast expression differences between deserted and reactive subTMEs (1,665 DEGs and 1,222 DEGs when adjusted for multiple comparisons (adj. DEGs) out of 5,708 total TME proteins), indicating profound regional proteomic differences within PDAC stroma. The subTME transcriptomes showed similar but less pronounced grouping in PCA (**Figure 2E**) and relatively fewer DEGs (3,607 DEGs and 1,091 adj. DEGs out of 17,945 TME transcripts). Since the histologically intermediate subTME phenotype (**Figure 1C**) grouped in between the reactive and deserted proteome samples by PCA (**Figure 2D**), showed intermediate differential protein expression (**Figure S2E**) and few DEGs when compared to reactive subTMEs (**Figure S2E**,**F**), we concluded it likely represents a transitory state and collapsed intermediate and reactive samples into one group.

We then integrated the patient-matched proteomic and transcriptomic datasets and compared DEGs between deserted and intermediary/reactive subTMEs in the intersection of 5,475 genes detected at both the RNA and protein levels. These were dominated by protein level changes with 1,612 DEGs and 1,194 adj. DEGs (**Figure 2F**), compared to 1,164 DEGs and 358 adj. DEGs at the RNA level (**Figure 2G**). Due to the small number of adjusted RNA DEGs, the intersection of adj. DEGs was small (n = 108) but expression changes were mostly reciprocated with 37 mutual downregulated genes, 58 mutual upregulated genes, and only 13 genes with discordant regulation (**Figure S2G, Table S2**). Notably, these subTME-specific 95 mutual transcript/protein adj. DEGs comprised several known players in tumor/stroma interaction (FBLN5, IGFBP5, IGFBP2) and markers of functionally distinct CAF subpopulations (ENG, LRRC15, DES, THBS4; **Figure 2H, Table S2**).

To pinpoint the biological functions unique to each PDAC subTME, we performed GSEA of the 1,612 integrated protein DEGs (**Figure 2F**), against 8,350 gene sets from the MSigDB v7.0 (biological pathways collection) and ConsensusPathDB. In line with their ECM-rich histological appearance (**Figure 1C**), deserted subTMEs exhibited a strong enrichment in ECM-related gene sets, ECM-related signaling terms, and humoral immune response pathways (classical complement pathway, B cell mediated immunity; **Figure 2I, Table S3**). Reactive subTMEs were enriched in gene sets related to cellular stress response mechanisms (heat shock, hypoxia, growth, metabolic stress), growth factor signaling with CAF-activating and immunomodulatory functions (FGF, EGF, TGF, PDGF, Wnt, GTPases), cytokine signaling (Interferons, TNF) and cellular immunity pathways (innate immunity, antigen presentation, TCR signaling; **Figure 2J, Table S4**). Thus, highly distinct gene expression profiles of deserted and reactive human PDAC subTMEs, especially at the protein level, indicated fundamental differences in stromal effector function.

### Reactive subTMEs harbor stromal cell communities with antitumor immune properties

To further dissect the functional biology and cellular composition of subTMEs, we constructed a deep tissue phenotyping platform for human PDAC tissues (**Figure 3A**). Briefly, in a tissue microarray (TMA) from PDAC resections (n = 165), we histopathologically determined the subTME present in each tissue core (n = 596) before applying a 25-marker (immuno)histochemistry panel and second harmonic generation (SHG) microscopy followed by semi-automated digital image analysis (**Methods**). Importantly, this allowed us to analyze the composition of subTMEs both across patients and in patient-paired reactive and deserted regions, while CK19-based estimation of TME and tumoral area and cell counts aided compartment-specific normalization (**Figure S3A**). SubTMEs indeed differed strongly in composition: reactive/intermediate subTMEs showed increased staining from markers of T cells (CD3), macrophages (CD68), endothelial cells (CD31), and CAFs (α SMA) while deserted subTMEs had high collagen content and maturation (**Figure 3B**), which was highly concordant with functional molecular annotation of subTME multiOMICs profiles (**Figure 2I,J**). Digital image analysis confirmed this differential expression pattern both globally across different patients and also, to a weaker extent, in patient-paired reactive and deserted regions (**Figure 3C**). Examination of bordering reactive and deserted regions in a small subset of tissue samples further illustrated the regional enrichment of these stromal components in the different subTME (**Figure 3D**). Of note, when we again singled out the intermediate subTMEs for separate comparison, they exhibited intermediate levels of these stromal components (**Figure S3B**), in line with their intermediate gene expression differences (**Figure 2D, S2E-F**).

**Figure 3.**
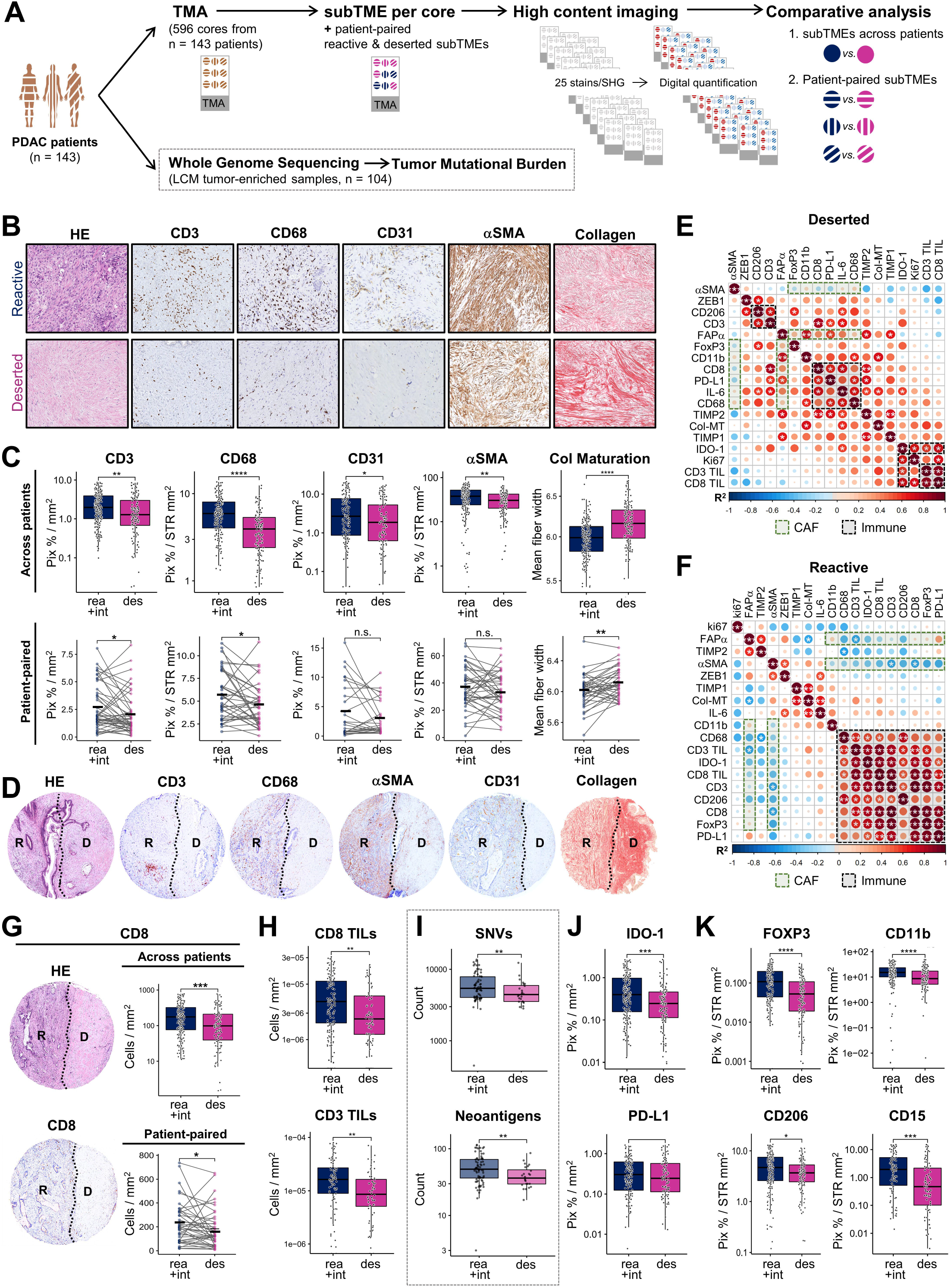
Reactive subTMEs harbour stromal cell communities with anti-tumor immune properties. See also **Figure S3** **A)** Schematic: Deep tissue phenotyping of subTMEs in 596 specimens from 143 PDAC patients. Adjacent sections of a tissue microarray (TMA) were stained with HE for subTME assessment of each core and analysed through stainings with Sirius Red, Masson’s Trichrome, and a 23-marker IHC panel or second harmonic generation (SHG) microscopy. Tumor mutational burden was analysed in a subset of 104 out of the 143 PDAC patients with matched whole genome sequencing (WGS). **B-D)** Reactive *vs*. deserted subTME regions stained for indicated markers. **B)** Representative images. **C)** Digital image analysis of IHC stains and collagen fibre width as assessed by SHG microscopy. Upper panel boxplots: all reactive vs. all deserted subTME regions; Wilcoxon test. Lower panel dot plots: patient-paired subTME regions; Wilcoxon signed-rank test. **D)** Examples of regions with bordering reactive and deserted subTMEs. **E-F)** Correlation matrix of selected cancer associated fibroblast (CAF) and immune markers in **E)** deserted subTMEs and **F)** reactive subTMEs. Spearman’s rank correlation method, paired. **G)** CD8 IHC staining. Representative images of bordering reactive and deserted subTMEs (*left*). Boxplot: all reactive vs. all deserted subTME regions. Wilcoxon test. Dot plot: patient-paired subTME regions. Digital image analysis of IHC stains. Wilcoxon signed-rank test (*lower panel*). **H)** CD8 and CD3 tumor infiltrating lymphocyte (TIL) counts in subTMEs. IHC manual counts; Wilcoxon test. **I)** Single nucleotide variant (*top*) and neoantigen counts (*bottom*) predicted from WGS profiles of resectable human PDACs (n = 104) compared by reactive *vs*. deserted main TMEs. Wilcoxon test. **J)** Checkpoint molecule expression in reactive *vs*. deserted subTMEs. IHC digital image analysis; Wilcoxon test. **K)** Immunosuppressive cell population markers in reactive *vs*. deserted subTMEs. IHC digital image analysis; Wilcoxon test. **C**,**E-F**,**G-K)** n.s. p>0.05, *p < 0.05, **p < 0.01, ***p < 0.001, ****p < 0.0001

To further define subTME-specific cell communities, we performed correlation analysis. Deserted subTMEs exhibited mild positive correlations of CAF markers FAPα and α SMA with immune markers and several small, scattered immune clusters (**Figure 3E**). In contrast, reactive TMEs showed a negative correlation of CAF markers with immune cells and an uninterrupted cluster comprising all immune cell and immunity markers (**Figure 3F**). Since GSEA analysis had also suggested prominent T cell-based immunity features in reactive subTMEs (**Figure 2J**), we further queried metrics of antitumor immunity. Reactive regions did show increased counts for total CD8 T cells (**Figure 3G**) and CD8^+^ and CD3^+^ tumor infiltrating lymphocytes (TILs) in direct contact with tumor cells (**Figure 3H**). We then assessed immunogenicity surrogates in 104 patient-matched tumor cell enriched whole genome sequencing profiles. Tumors with a reactive-dominant TME exhibited higher SNV and neoantigen counts (**Figure 3I**) and even though the global genomic analysis could not account for regional differences, this data altogether indicated a hot, T cell inflamed immune phenotype in reactive subTMEs. Indeed, reactive subTMEs also had higher IDO-1 levels and a trend to increased PD-L1 levels (**Figure 3J**), along with increased markers for suppressive immune cell types including FOXP3 (regulatory T cells), CD11b and CD15 (MDSCs), and CD206 (M2 macrophages; **Figure 3K**), suggesting a cell-based regulatory response typical for PDAC (Fan et al., 2020; Steele et al., 2020). Intratumoral subTMEs thus differed widely in composition and likely harbor distinct immune milieus.

### CAF communities in human PDAC subTMEs establish coordinated phenotypes with distinct behavior and function

Given their central role in the histological appearance of subTMEs (**Figure 1C**) and known immunomodulatory functions, we next queried the subTME-specific CAF compartment. The subTME multiOMIC datasets exhibited distinct CAF marker profiles (**Figure 4A**), such that different CAF subpopulations might preferentially localize to deserted or reactive regions. Yet, due to the different cellularity of subTMEs (**Figure 4B**), we could not definitively discriminate whether these marker differences stemmed from differential CAF numbers and/or differential per-cell expression levels. We thus extracted CAFs from 13 fresh PDAC specimens with histopathologically confirmed subTMEs (**Figure 4C**) for single cell and functional characterization. Remarkably, the growth patterns of subTME CAF culture monolayers closely recapitulated the characteristic histology of their originating subTMEs (**Figure 4D**). To test whether these distinct phenotypes were accompanied by behavioral differences we focused on migration and proliferation, two disparate fibroblast functionalities (De Donatis et al., 2010). Reactive subTME CAF cultures were more motile (**Figure 4E**) whereas deserted subTME CAF cultures grew faster (**Figure 4F**). Thus, human PDAC subTMEs harbored phenotypically and behaviorally distinct CAFs.

**Figure 4.**
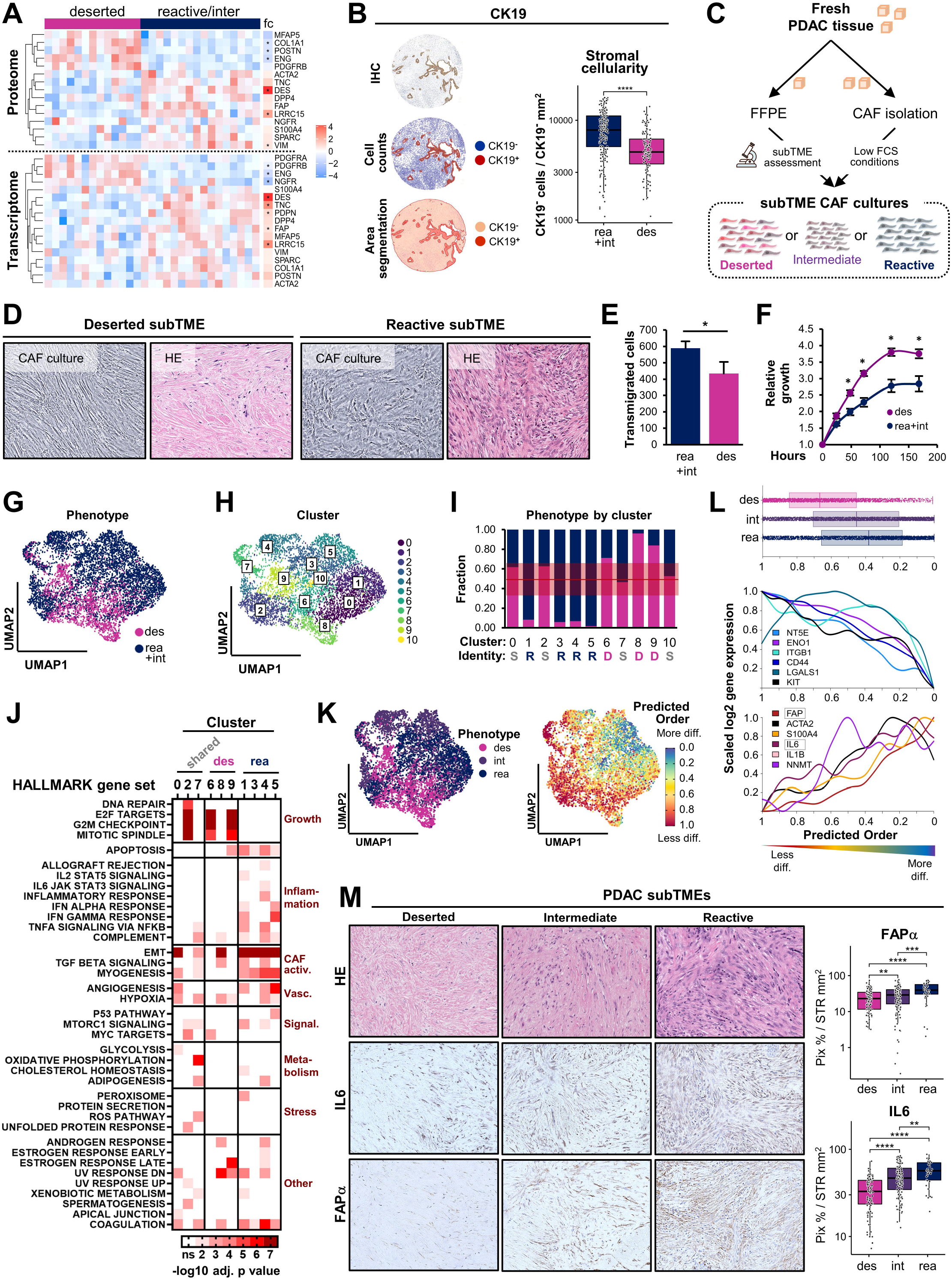
CAF communities in human PDAC subTMEs establish coordinated phenotypes with distinct behaviour and function. See also **Figure S4** **A)** Heatmap: Selected CAF markers in multiOMIC profiles from reactive and deserted subTMEs. **B)** Estimation of TME cellularity through CK19 IHC. Representative images (*left*). Stromal cell count by stromal area in reactive *vs*. deserted subTMEs (right). Wilcoxon test. **C)** Schematic: Isolation of subTME CAFs from fresh human PDAC tissues (n = 13). **D)** Morphological comparison of subTME *ex vivo* CAF cultures at confluency (phase contrast) and originating subTME tissue histology (HE stain). **E)** Transwell migration of reactive (n = 4) *vs*. deserted (n = 3) subTME CAF cultures towards 10% FCS. Mean + SEM. Wilcoxon test. **F)** Relative in vitro growth of reactive (n = 4) *vs*. deserted subTME CAF cultures (n = 3). Mean +/-SEM. Wilcoxon test. **G)** UMAP embedding of single cell transcriptomes of 6,331 CAFs originating from deserted and reactive/intermediate subTMEs of 10 PDAC patients. **H)** UMPA embedding overlay showing 11 clusters. **I)** Breakdown of CAF originating subTME to each cluster. Cluster identity assigned when >66% contribution of one subTME. S, shared; R; reactive; D, deserted. **J)** Heatmap summarizing HALLMARK gene sets (rows) upregulated in each cluster (columns). Clusters grouped by subTME identify from (**I**) **K)** UMPA embedding overlays showing cells from deserted (’des’), reactive (‘rea’) and intermediate (‘int’) subTMEs as separate groups (*left*) and CytoTRACE scores (*right*) reflecting predicted order of differentiation from values 1 (less diff, red) to 0 (most diff, blue). **L)** CytoTRACE scores of CAFs grouped by originating subTME (*top*). Selected pluripotency markers (*middle*) and CAF activation markers (*bottom*) displayed as a smoothing spline over the moving average of 200 cells. **M)** IL6 and FAPα IHC in human PDAC subTME tissues. Representative images (*left*). Boxplots: digital IHC stain quantification comparing subTMEs; Wilcoxon test. **B**,**E**,**F**,**M)** n.s. p>0.05, *p < 0.05, **p < 0.01, ***p < 0.001,****p < 0.0001

To query the CAF subpopulations in subTME CAF cultures, we performed single cell RNAseq on 10 in vitro cultures. 6,331 CAFs with a minimum of 200 genes detected per cell and <3% mitochondrial genes were carried into further analyses (**Figure S4A**). Unsupervised graph-based clustering showed that CAFs largely grouped by their originating subTME (**Figure 4G**) yet comprised a total of 11 individual clusters (**Figure 4H**). All 11 clusters were found in all 10 patients, albeit at varying levels (**Figure S4B**). Clusters 1/3/4/5 were enriched (>66%) in reactive subTME-derived CAFs, clusters 6/8/9 were enriched in deserted subTME-derived CAFs, and clusters 0/2/7/10 were shared among both groups at 33-66% frequency each (**Figure 4I**). Altogether, subTME CAF isolates contained several clusters, some of which were shared, and others were subTME-specific. We next analyzed whether these clusters represent known inflammatory CAF (iCAF) and myofibroblastic CAF (myCAF) subpopulations (Elyada et al., 2019; Kieffer et al., 2020). There was no overlap with shared clusters 2/7 but shared cluster 0 aligned with all myCAF signatures and deserted-enriched clusters 8/9 with the pan-iCAF signature (Elyada et al., 2019) (**Figure S4C**). All reactive-type CAF clusters showed enrichment in several iCAF and myCAF signatures, including ecm-myCAF, TGFb-myCAF, and wound-myCAF, shown to be immunomodulatory and predictive of anti-PD-1 response in other cancers (Kieffer et al., 2020). Overall, only few clusters in the subTME CAF isolates resembled known iCAF *vs*. myCAF subpopulation differences.

We proceeded to functional molecular annotation of each cluster through hypergeometric testing of DEGs (**Figure S4D**) against the HALLMARK gene sets (Liberzon et al., 2015). Deserted CAF clusters 6/9 showed strong enrichment of growth-related HALLMARK terms (**Figure 4J**); in line with an increased expression of G2M and S phase gene signatures (**Figure S4E**) and the increased growth rates of deserted-type CAF cultures (**Figure 4F**). Alternatively, reactive subTME-derived CAFs exhibited broader functional molecular properties: the inflammation-related HALLMARK gene sets (allograft rejection, IL2-STAT5, IL6-JAK-STAT, inflammatory response, interferons, TNFalpha, complement) were thoroughly enriched across 3 out of 4 reactive-type clusters (**Figure 4J**), suggesting that CAFs can contribute to the prominent inflammatory features of the reactive subTMEs we observed in tissues (**Figure 2J, Figure 3C-K**). Furthermore, all reactive-type CAF clusters exhibited strong enrichment of the EMT gene set (**Figure 4J**), likely reflective of the increased motility of reactive subTME CAFs (**Figure 4E**). Myogenesis and TGFβ gene sets indicative of increased CAF activation status were also enriched across reactive CAF clusters, as were apoptosis and vascularization-related terms (**Figure 4J**). Since the multi-subpopulation CAF communities in deserted and reactive subTME cultures appeared to be in ‘coordinated states’, represented by cluster-overarching functional profiles (**Figure 4J**) and distinct morpho-histological (**Figure 4D**) and behavioral (**Figure 4E**,**F**) phenotypes, we asked whether this was a discrete or continuous phenomenon. We singled out the intermediate subTME CAFs into a separate group and inferred a differentiation trajectory through CytoTRACE (Gulati et al., 2020). This predicted an order of differentiation states from deserted-type CAFs (less differentiated) to reactive-type CAFs (more differentiated) that indeed transitioned through the intermediate-type CAFs (**Figure 4K**, red to yellow to blue). On this deserted-intermediate-reactive trajectory, the moving average of pluripotency markers (NT5E, ENO1, ITGB1, CD44, LGALS1, KIT) consistently decreased whereas CAF activation markers (FAPα, ACTA2/α SMA, S100A4), regulators (NNMT), and inflammatory phenotype markers (IL6, IL1B) increased (**Figure 4L**). Similarly, in tissues, CAF activation maker FAPα and inflammatory phenotype marker IL6 increased from deserted to intermediate to reactive subTME regions (**Figure 4M**). Altogether, this suggests that subTMEs comprise complex CAF communities in different differentiation states with coordinated phenotypes and distinct behavior and function.

### Reactive subTMEs support a proliferative, basal-like, and poorly differentiated tumor cell phenotype

To investigate the relationship between subTMEs and PDAC progression, we next focused on the multiOMICs profiles of tumor epithelium extracted from reactive vs. deserted subTMEs (**Figure 2A**). These exhibited 492 DEGs and 9 adj. DEGs at the protein level and 1,223 DEGs and 0 adj. DEGs at the RNA level (**Figure S5**). In contrast to the above proteome-dominated differences in TME fractions (**Figure 2F,G**), the PDAC malignant epithelium showed a comparable extent of differential gene expression at the integrated RNA and protein levels (proteome: 465 DEGs and 9 adj. DEGs; transcriptome: 368 DEGs and 0 adj. DEGs; **Figure 5A**). In GSEA using HALLMARK gene sets, malignant epithelium from reactive subTMEs was enriched in cell cycle progression (G2M phase, E2F targets), oncogenic signaling (MYC), and interferon response terms. Malignant epithelium extracted from deserted subTMEs showed enrichment in cellular metabolism (fatty acids, oxidative phosphorylation), protein secretion, and drug metabolism terms (**Figure 5B**). We thus next functionally tested these putative effects of subTME CAFs on PDAC biology using patient-derived organoids (PDOs) and validated findings in our tissue imaging and clinical datasets (**Figure 5C**).

**Figure 5.**
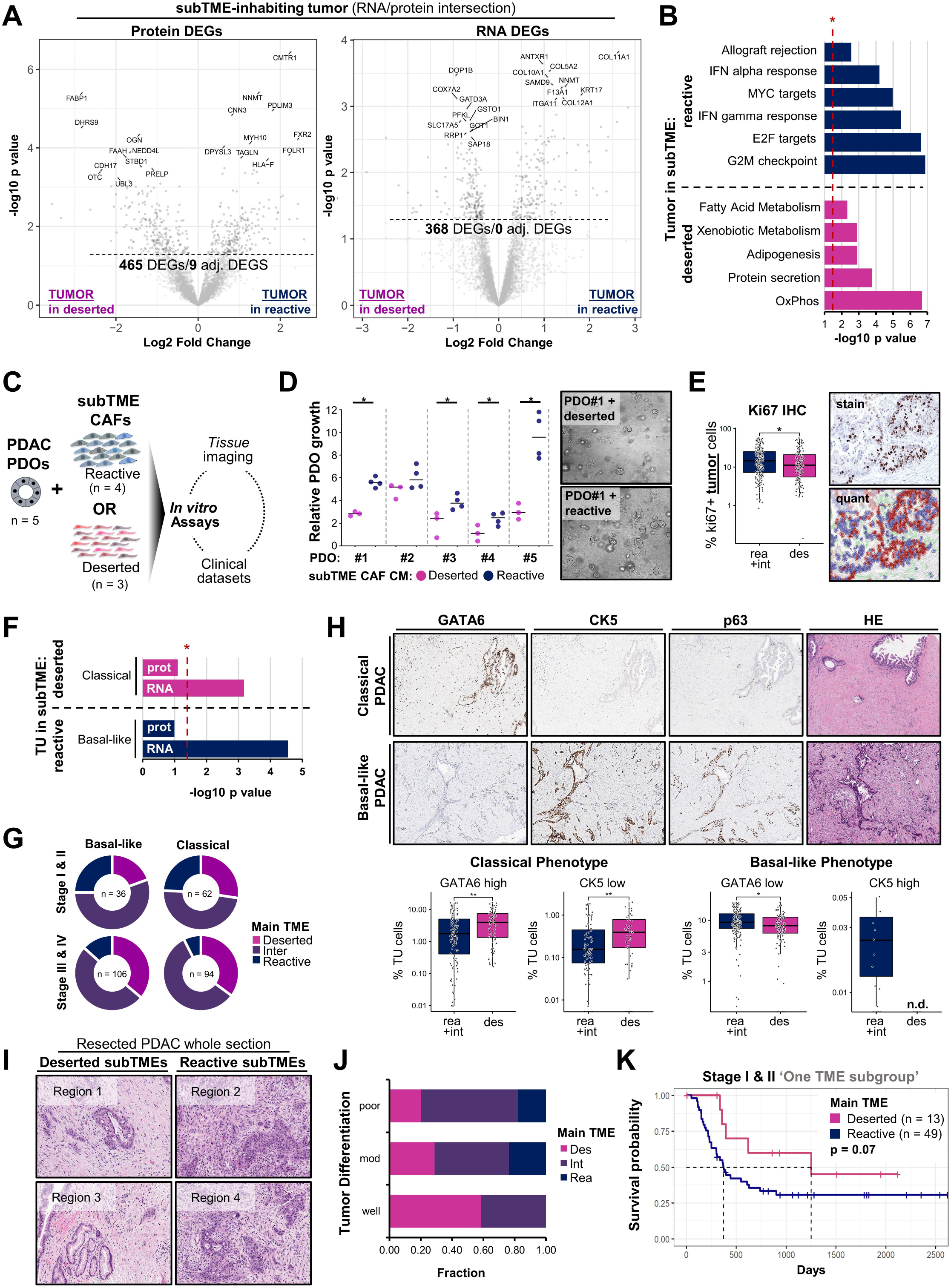
Reactive subTMEs support a proliferative, basal-like, and poorly differentiated tumor cell phenotype. See also **Figure S5** **A)** Volcano plots: DEGs in tumor epithelium extracted from deserted vs. reactive subTMEs at protein (*left*) and RNA level (*right*) following proteome-transcriptome integration (n=29). **B)** Bar plot summarizing enrichment p values of HALLMARK gene sets up-regulated in tumor epithelium extracted from deserted *vs*. reactive subTMEs. Wilcoxon test. Dashed red line = significance threshold. **C)** Schematic: Approach to interrogation of subTME effects on human PDAC utilizing patient-derived organoids (PDOs), tissue imaging, and clinical outcome data. **D)** Growth of human PDAC patient-derived organoids (PDOs; n = 5) in conditioned media (CM) from reactive (n = 4) *vs*. deserted (n = 3) subTME CAFs. 7-day endpoint normalized to 12-h seeding reference, Wilcoxon test. **E)** ki67 IHC in 596 specimens from 143 PDAC patients. Boxplots: ki67-positive fraction of tumor cells in reactive *vs*. deserted regions; Wilcoxon test (*left*). Stain and detection classifier for tumor cell-specific quantification (*right*). **F)** Bar plot summarizing enrichment p values of basal-like/classical gene sets in tumor epithelium extracted from deserted or reactive subTMEs. Wilcoxon test. Dashed red line = significance threshold. **G)** Frequencies of main TME phenotypes in basal-like *vs*. classical PDAC cases in Toronto in-house cohorts. **H)** IHC analysis of basal-like/classical markers in tumor cells in different subTMEs in 596 specimens from 143 PDAC patients. Representative images (*top*). IHC image analysis (*bottom*); n.d., not detected. Wilcoxon test. **I)** Regional association of varying tumor differentiation with different subTMEs. Shown are 4 regions from one representative tumor. **J)** Global association of tumor differentiation with main subTME phenotype in human PDAC (n = 143) **K)** Kaplan-Meier analysis of disease-free survival for patient subset with one main TME phenotype. Resection cohort; strata: main TME phenotype; Log rank test. **A**,**B**,**D**,**E**,**F**,**H)** *p < 0.05, **p < 0.01, ***p < 0.001,****p < 0.0001

As suggested by GSEA (**Figure 5B**), PDAC PDOs proliferated more in conditioned media from reactive-type CAFs (**Figure 5D**). Likewise, in tissues, tumor cells located within reactive subTMEs exhibited a higher proliferative index (**Figure 5E**). Thus, reactive subTMEs, and particularly the CAFs within these regions, can support tumor cell growth. Next, we followed up on the enrichment of the MYC hallmark gene set in reactive subTME-derived tumor cells (**Figure 5B**), since MYC activity is a central hub in the establishment of the aggressive squamous/basal PDAC subtype (Bailey et al., 2016; Hayashi et al., 2020). The basal-like gene signature (Moffitt et al., 2015) was enriched on RNA and on protein levels in malignant epithelium extracted from reactive subTMEs (**Figure 5F**). Globally, squamous/basal-like cases showed a lower frequency of a deserted-dominant TME in PDAC resections (n = 98) and a higher frequency of reactive-dominant TME in stage III&IV cases (n = 200; **Figure 5G**). Likewise, tumor cells located in reactive subTMEs showed a more basal-like phenotype profile by differential expression of squamous marker CK5 and differentiation factor GATA6 (**Figure 5H**). Moreover, reactive and deserted subTMEs within the same tumor were often accompanied by regional changes in tumor phenotypes i.e., reactive subTMEs frequently harbored tumor cells with squamous features, solid sheets or cribriform glands while neighboring deserted subTMEs were often inhabited by well-polarized glands (**Figure 5I**), further reflected in a strong association of tumor differentiation with frequency of deserted-dominant TMEs (**Figure 5J**). Accordingly, in resectable patient subgroups with a single dominating TME phenotypes TME type (**Figure 1K**, strata in grey), a reactive-dominant TME did trend towards a shortened disease-free survival (**Figure 5K**). Altogether, the deserted subTME appeared to support tumor differentiation while a reactive subTME promoted tumor progression through a proliferative, de-differentiated, squamous/basal-like tumor cell phenotype.

### The deserted subTME is chemoprotective

Given the prominent role proposed for stroma in hampering PDAC treatment response (Liang et al., 2017), we next investigated the effects of subTMEs on chemoresistance. Conditioned media from deserted-type CAFs increased Gemcitabine resistance in 4 out of 5 PDAC PDOs (**Figure 6A**), compared to conditioned media from reactive-type CAFs. Furthermore, in patients from the COMPASS trial (O’Kane et al., 2020), a deserted-dominant TME was strongly associated with poor response (**Figure 6B**) and reduced average tumor size change (**Figure 6C**) during 1^st^ line chemotherapy. Further, deserted-dominant TME frequency was lowest in partial responders and increased stepwise to patients with stable and with progressive disease (**Figure 6D**). Consequently, a deserted-dominant TME was a clear negative prognostic indicator in stage III, IV patients (**Figure 6E**), altogether suggesting that deserted subTMEs negatively affect late-stage outcome through chemoprotective effects. We next asked whether chemotherapy can affect the TME phenotype. Neoadjuvant treated PDACs (n = 23) frequently exhibited a deserted-dominant TME, compared to treatment-naïve resections (n = 143; **Figure 6F**), and this was accompanied by lower stromal immunoreactivity for IL6 and FAPα (**Figure 6G**), indicative of a deserted-like CAF phenotype (**Figure 4L-M**). We then evaluated 15 pairs of patient-matched pre-treatment/post-treatment biopsies (**Figure 6H**) and observed switches from a reactive or intermediate-dominant to a deserted-dominant phenotype in 9/15 cases and 1/15 reactive-to-intermediate switch; 1/15 remained deserted and 3/15 remained intermediate. One case exhibited a ‘reverse’ switch (deserted to intermediate), accompanied by a switch in epithelial phenotype from classical to basal-like also. Taken together, deserted subTMEs became more frequent upon chemotherapy and had a chemoprotective effect, at least in part mediated by CAFs. Thus, chemoprotection and disease promotion appear be independent stromal support functions, executed by deserted and reactive subTMEs, respectively.

**Figure 6.**
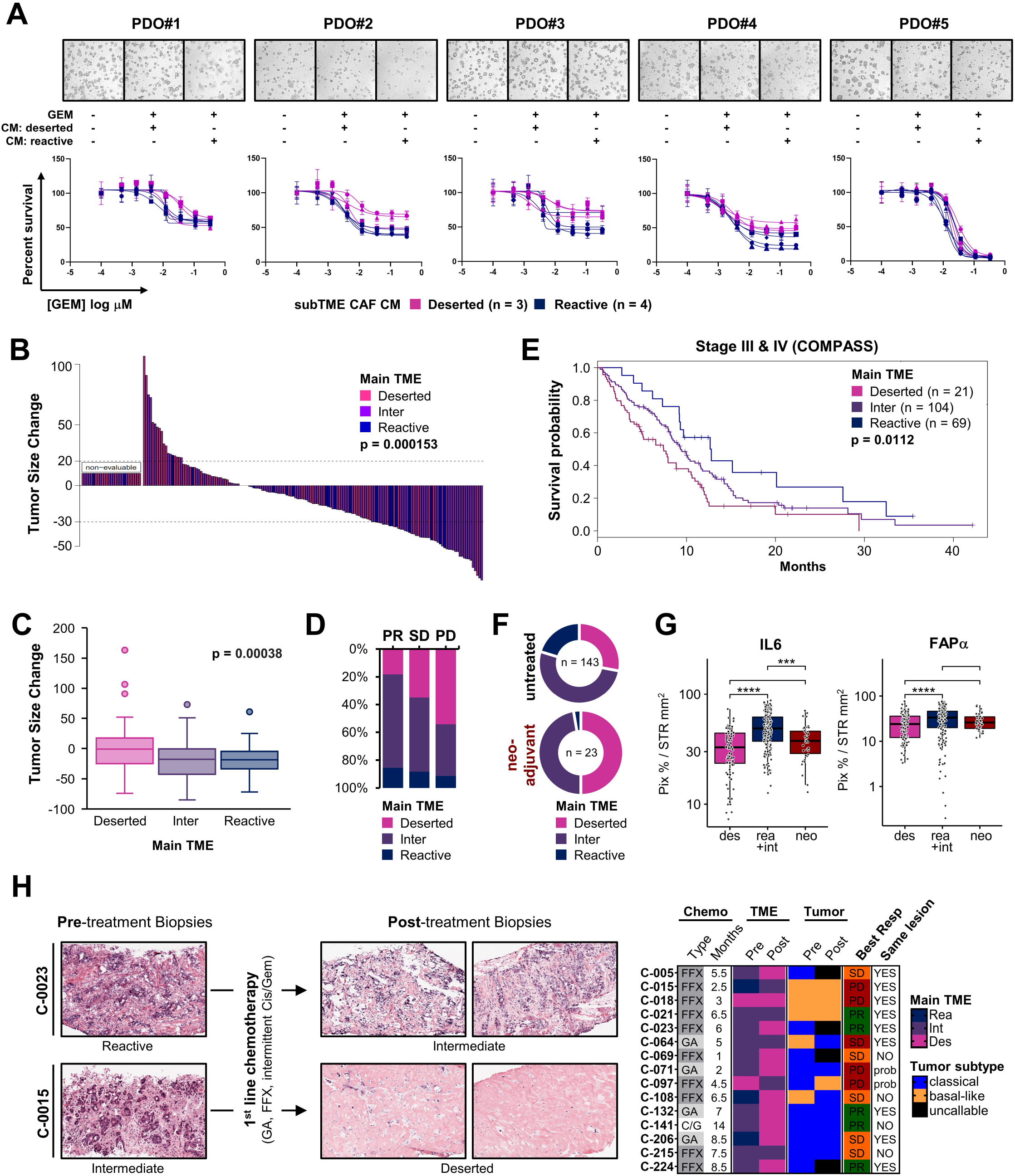
The deserted stroma phenotype is chemo-protective. **A)** Gemcitabine sensitivity of human PDAC patient-derived organoids (PDOs; n = 5) in conditioned media (CM) from reactive (n = 4) *vs*. deserted (n = 3) subTME CAFs. Upper panel: brightfield images of representative wells. Lower panel: dose response assays. **B-D)** Response to 1^st^ line chemotherapy by main TME phenotype in in-house COMPASS cohort. **B)** Waterfall plot: tumor size change; Fisher’s exact test. **C)** Average tumor size change; Kruskal-Wallis test. **D)** Best response. PD, progressive disease; SD, stable disease; PR, partial response. **E)** Kaplan-Meier analysis of overall survival. COMPASS cohort; strata: main TME phenotype; Log rank test. **F)** Main TME phenotypes frequencies in PDAC resections with and without neoadjuvant treatment. **G)** IL6 and FAP expression in subTMEs *vs*. neoadjuvant treated PDAC tissues. IHC digital image analysis; Wilcoxon test; ****p < 0.0001 **H)** Switch of main TME phenotypes during chemotherapy. Representative images from two progression biopsies (*left*). Heatmap summarizing pre-/post-treatment pairs (n = 15; *right*). FFX, folfirinox; GA, gemcitabine plus (nab)-paclitaxel; C/G, intermittent cisplatin /gemcitabine; PD, progressive disease; SD, stable disease; PR, partial response; prob, likely same lesion based on records but not specifically confirmed.

### Prediction of subTMEs from bulk RNAseq profiles corroborates prognostic impact of stromal heterogeneity

Finally, we examined whether we could leverage the subTME-specific expression differences to infer TME phenotypes and heterogeneity from public bulk RNAseq profiles (**Figure 7A**). A Random Forest machine learning model was trained on reactive *vs*. deserted subTME transcriptomes (n = 23). In a feature selection step, 72 genes were selected by their impact score in the model and the overall impact score distribution (**Figure S6A**). The resulting ‘TME PHENOtyper’ model performed with >95% accuracy at a virtual tumor background of 50% (**Figure 7B**) and was then applied to 177 bulk tissue RNAseq profiles from the TCGA PAAD dataset (**Figure 7C**). TME PHENOtyper predicted the dominant TME phenotype with a performance of 93.4% in TCGA samples with >50% stroma (**Figure 7D**) and an accuracy of 74.8%, as benchmarked against histopathological review. SubTME scores did not correlate with ABSOLUTE-based (Carter et al., 2012) tumor cellularity estimates (**Figure S6B**) further indicating that the model is robust even with varying tumor content in bulk samples. Finally, we tested whether TME PHENOtyper could be used to identify the prognostic subgroups of patients with one main TME phenotype vs. co-occurring subTMEs (**Figure 1K**) by using subTME score quantiles as a proxy TME heterogeneity (**Figure 7E**). Indeed, high subTME scores (top 25% reactive or deserted score) yielded better outcome compared to middle quantiles with moderate reactive and deserted scores (**Figure 7F**) and separation of survival curves increased when we repeated the same analysis with top 12.5% quantiles (**Figure 7G**), suggesting that increased TME heterogeneity as a function of subTME co-occurrence is continuously associated with poor disease outcome.

**Figure 7.**
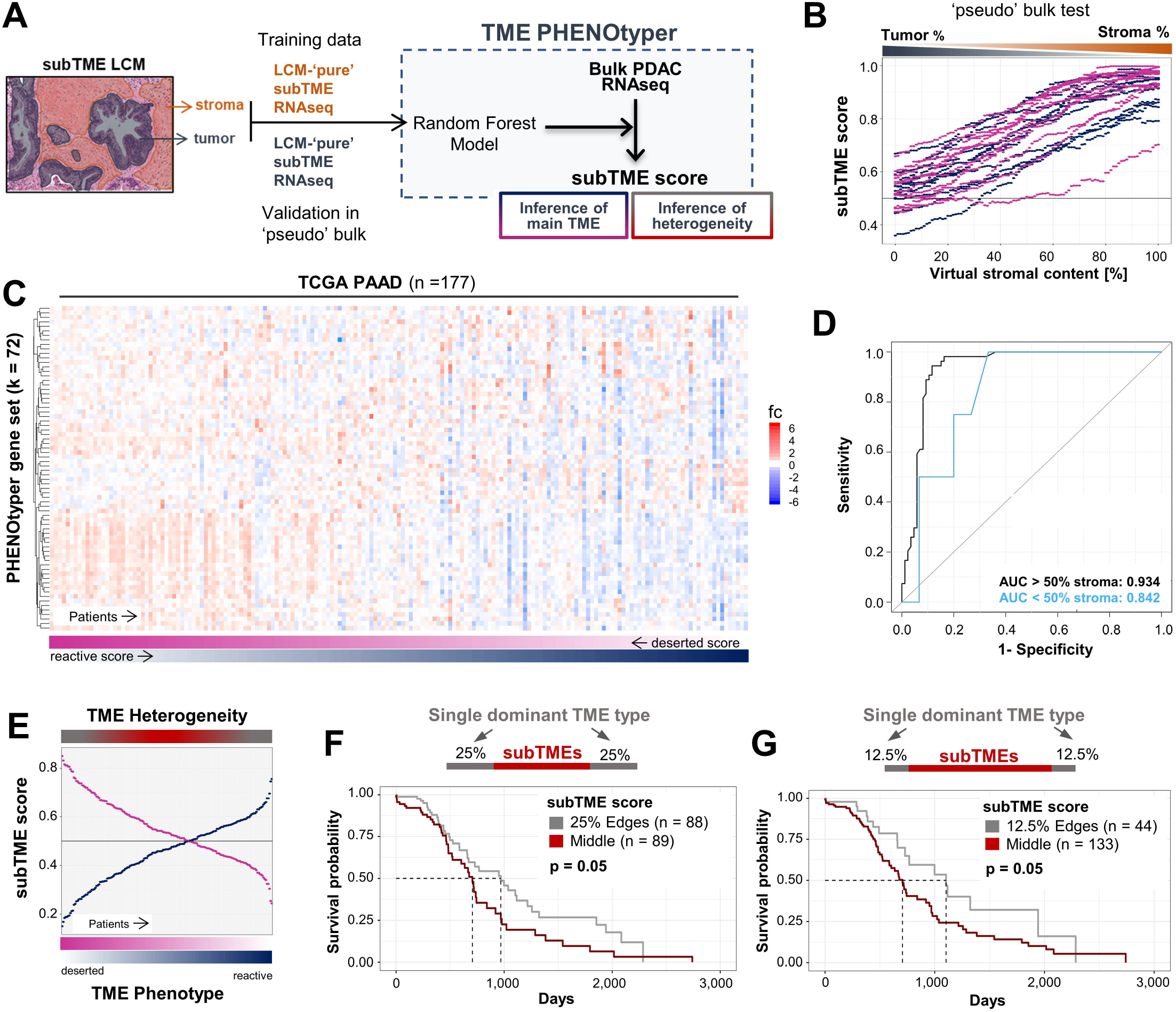
Prediction of stromal subTMEs from bulk RNAseq profiles corroborates prognosti impact of stromal heterogeneity. See also **Figure S6** and **Tables S5-S6**. **A)** Schematic: Workflow of training TME PHENOtyper, a Random Forest model for prediction of subTMEs from bulk RNAseq samples. **B)** Robustness of TME PHENOtyper subTME scores against incremental virtual mixing of patient-matched tumor and TME RNAseq profiles. **C)** Heatmap visualizing expression of TME PHENOtyper 72-gene set in TCGA PAAD cohort. **D)** Validation analysis of subTME predictions by TME PHENOtyper in cases with >50% stroma (black line) and <50% stroma (blue line) in TCGA PAAD bulk RNAseq data. **E)** Inference of stromal heterogeneity from middle quantiles of subTME scores. **F-G)** Kaplan-Meier analysis of overall survival. TCGA PAAD cohort. Strata: **F)** combined top 25% reactive and deserted subTME scores *vs*. rest; **G)** combined top 12.5% reactive and deserted subTME scores *vs*. rest. Log rank test.

## Discussion

Regional heterogeneity is a long-standing obstacle to defining precise stromal contributions to PDAC progression and chemoresistance. This study systematically integrates clinical histopathology with multi-tiered OMICs analyses and reveals how stromal phenotypic heterogeneity in PDAC is a function of regional TME programs, which cooperate to actively drive PDAC progression. Specifically, we discovered two overarching microenvironmental states with distinct tumor-promoting and chemo-protective roles in human PDAC. The underlying reactive and deserted sub-microenvironments (subTMEs) demonstrated regional variance in TME composition, phenotype, and function. Namely, reactive subTMEs were vascularized, immune-hot, and tumor-promoting but appeared more sensitive to chemotherapy. They frequently co-existed intratumorally with deserted subTMEs which supported tumor differentiation but were chemoprotective *in vitro* and associated with poor response to treatment in patients. Accordingly, tumors benefited from the concomitant presence of reactive and deserted subTMEs, which resulted in poor outcome for patients with a phenotypically heterogenous TME.

CAFs are a central element in establishing subTMEs. We unexpectedly observed up to 10 subpopulations in subTME CAF cultures that were unified in their subTME-specific differentiation states and together established coordinated phenotypes with distinct behavior and function. SubTMEs thus represent organizational units of complex CAF communities, likely much more nuanced in tissues than in our *in vitro* cultures. We observed partial overlap with original (Elyada et al., 2019) and recently refined iCAF/myCAF subpopulations (Kieffer et al., 2020) but also found additional subpopulations along the continuum of CAF differentiation states from deserted to intermediary to reactive. Noticeably, this CAF differentiation potential was associated with distinct tumor-related functions and, similar to stem cells (Gulati et al., 2020), was marked by RNA diversity and pluripotency markers. In sum, stromal heterogeneity in PDAC appears to result largely from transitions between two CAF cell states that regionally govern multifaceted CAF and immune cell communities. Consequently, the PDAC TME, despite its complexity and the myriad possible combinations of its inhabiting cell populations, may de facto self-organize into relatively few states of functional relevance. While our current understanding of the processes governing tissue self-organization and patterning is very limited (Muthuswamy, 2021; Santos and Liberali, 2019), the conservation of the here-identified PDAC subTME histological patterns across stages and disease sites does strongly suggest that they are not random but reflect a definable and reproducible repertoire of tissue responses to chronic injury.

Stromal heterogeneity, as defined by the co-occurrence of reactive and deserted subTMEs within a single patient, was strongly linked to poor outcome. Our findings suggest that tumors with such a heterogenous TME have concomitant access to tumor-promoting and chemo-protective stromal elements so that the associated poor survival likely results from complementary effects of subTMEs on tumor pathobiology. Our compartment-specific approach revealed that reactive subTME and the basal-like/squamous subtype of tumor cells were partially linked in a regional and a global manner, which is especially significant in light of recent findings that the transcriptional epithelial subtypes, like subTMEs, can co-exist in the same tumor (Chan-Seng-Yue et al., 2020; Hayashi et al., 2020). These findings here and of others (Ligorio et al., 2019) show that the TME can promote heterogeneity of tumor cell phenotypes. Moreover, subTMEs instigated regional immune heterogeneity and reactive subTMEs presented as distinct immune-hot regions. This corroborates known links between squamous differentiation and tissue inflammation (Somerville et al., 2020), altogether tying regional TME phenotypes to both epithelial and immune heterogeneity in human PDAC. The subTME-anchored regional immune heterogeneity in human PDAC has implications for immunotherapy design and patient selection. Beyond that, a chemotherapy-induced TME switch towards the poorly vascularized, ECM-rich and chemoprotective deserted phenotype in patient-paired progression biopsies indicates its potential role in acquired chemoresistance.

Altogether, this study reveals that variations in spatial phenotypic TME features mark transitions between two fundamental microenvironmental states in PDAC, each with its own cellular and molecular underpinnings, that produce patient-specific stromal heterogeneity and are tightly linked to clinical outcomes. We demonstrate how incorporating this regional TME concept into standard histopathology not only complements but radically streamlines the extraction of knowledge from deep molecular profiling, immune-phenotyping, and patient outcome data. This opens a new dimension for understanding the tumor-stroma-immune interplay in PDAC progression and treatment response, along with a new and highly accessible avenue to TME-cognisant tumor classification, prognostication, and prediction of therapeutic response for PDAC patients.

## Acknowledgments

We thank Paul Waterhouse for detailed review of this manuscript and all members of Khokha lab for helpful discussions. This work was supported by grants to RK from the Canadian Cancer Society Research Institute (CCSRI). BTG held fellowships from the Princess Margaret foundation, EMBO (ALTF 116-2018) and the Alexander von Humboldt Foundation (DEU 1199182 FLF-P). CWM held a Banting Postdoctoral Fellowship. This study furthermore supported by the Ontario Institute for Cancer Research (PanCuRx Translational Research Initiative) through funding provided by the Government of Ontario, the Wallace McCain Centre for Pancreatic Cancer supported by the Princess Margaret Cancer Foundation, the Terry Fox Research Institute, the Canadian Cancer Society Research Institute, and the Pancreatic Cancer Canada Foundation. SG is the recipient of an Investigator Award from OICR. Additional support was provided by the Deutsche Forschungsgemeinschaft (DFG) SFB 850 subprojects C9 (to AD, MB), Z1 (to MB), and from the German Federal Ministry of Education and Research (BMBF) by MIRACUM within the Medical Informatics Funding Scheme (FKZ 01ZZ1801B, to MB). We thank Dr. Zev Gartner for generously providing MULTIseq reagents. We acknowledge the contributions of team members in the Diagnostic Development platform and the Genomics Program (genomics.oicr.on.ca) at OICR.

## Author Contributions

BTG conceived the study and designed and performed most of the experiments and data analyses. AD, GA, GHJ, RD, KA, FV, CWM, AM, JMR, PrB, PBr, GOK, JW, JK, LT, NR contributed to experiments, data acquisition and/or analysis. MB oversaw bioinformatics analyses. SF performed histopathological review. MB, SG, TK, and RK supervised the study. BTG wrote the manuscript which other authors edited and approved.

## Declaration of Interests

The authors declare no competing interests.

## Materials & Methods

### Cohort Description

The study includes 391 specimens from 376 individuals with histologic diagnosis of PDAC (143 resectable tumor specimens from treatment-naïve patients with stage I/II PDAC; 23 resectable tumor specimens from neoadjuvant-treated PDAC patients; 210 diagnostic biopsy specimens and 15 patient-paired longitudinal biopsies from 210 stage III/IV PDAC patients). A summary of the patient cohort is provided in **Table S1**. All patients provided written informed consent allowing the molecular characterization of their tumor samples and follow-up on their clinical information. Resectable tumors were obtained from the UHN Biospecimens Program, and advanced tumors were obtained from the COMPASS trial (no. NCT02750657). Patient samples were mostly accrued at Princess Margaret Cancer Centre at the University Health Network (Toronto, Canada) and reported in previous studies (Aung et al., 2018; Connor et al., 2019; O’Kane et al., 2020). All studies were approved by University Health Network Research Ethics Board (Nos. 03-0049, 08-0767, 15-9596, 17-6106, 16-5380). The study complies with all relevant ethical regulations.

### TCGA data retrieval

The validation cohort consisted of 177 PDAC patients from The Cancer Genome Atlas (TCGA)(TCGA Research Network, 2017) data portal. The TCGA database was accessed on February 3rd, 2020 via the TCGAbiolinks package (Colaprico et al., 2016). 182 PAAD RNAseq V2 data sets were downloaded for analyses. Five samples were filtered out due to annotation of a non-pancreas primary site.

### Histopathological Review

Haematoxylin and eosin (H&E) stained slides from all patients were reviewed by a pathologist (SF). All available tumor-containing whole sections (resectable PDAC cases) or biopsy cores (advanced PDAC cases) per patient were semi-quantitatively assessed for the presence of specific, reproducible morphological patterns. This identified 1. a ‘deserted’ subTME, defined as paucicellular region with thin, spindle-shaped fibroblasts embedded in loose extracellular matrix and fully matured collagen fibers, often with keloid and myxoid features; 2. a ‘reactive’ subTME, defined as region containing fibroblasts with plump morphology and enlarged nuclei, little acellular components but often a rich inflammatory infiltrate; 3. regions with intermediate levels of these features, which were recorded as ‘intermediate’ subTMEs. All subTMEs present were recorded and the subTME with >50% estimated area contribution was assigned as main TME phenotype per case. Tumor differentiation was assessed from the same specimens using the standard pathological grading scheme into either well differentiated, moderately differentiated, and poorly differentiated based on the lowest differentiation grade observed. HE slide scans from diagnostic specimen of 159 cases for the TCGA validation cohort were obtained from https://portal.gdc.cancer.gov/ and assessed identically.

### Laser capture microdissection (LCM)

LCM of freshly frozen tissue samples was performed on a Leica LMD 7000 instrument as previously described (Connor et al., 2019). Briefly, frozen tissue maintained in vapor-phase liquid nitrogen was embedded in OCT cutting medium and sectioned in a cryotome into 10-µm thick sections. 9-14 sections per case were mounted on PEN membrane slides (Leica) and lightly stained with haematoxylin to facilitate microscopic identification of tumor and stroma areas. An adjacent section per case was haematoxylin-eosin (HE) stained and used in parallel at a light microscope to aid histologic confirmation of stromal and tumoral features. LCM was performed within 48 h after sections were cut to minimize nucleic acid and protein degradation.

First, microdissected tumor epithelium from defined subTME regions was collected by gravity into the caps of sterile, RNAse-free microcentrifuge tubes. Approximately 50,000-100,000 tumor cells were collected for each RNA and protein extraction sample. Next, any non-TME components (e.g. residual acini and normal ducts, large blood vessels, nerves, scattered tumor nests) were microdissected from the TME and discarded. Finally, approximately 50,000 cells from thereby cleaned subTME-specific stromal regions were collected for each RNA and protein extraction samples. Arcturus PicoPure Extraction Buffer was added directly to RNA extraction samples, while protein extraction samples were spun down and left dry. All samples were then stored at −80°C until further processing.

### Shotgun Proteomics

Shotgun proteomics on dissected fractions from OCT embedded tissue has been described previously (Sinha et al., 2019). Removal of the OCT compound was performed using various dilutions of ethanol with water as follows. At each step, the tubes were centrifuged at 13,200 g for 3 min, and the supernatant was discarded. Initially, 1 mL of 70% (v/v) ethanol was added to each conical tube, followed by 30 s of vortexing. Subsequently, tissue pellets were resuspended with 100 µL of 100% ethanol, 70% ethanol, 85% ethanol and 100% ethanol with an additional 5 min incubation at room temperature and 30 sec vortexing. Finally, pellets were washed with PBS twice, spun down, snap frozen in liquid nitrogen and stored at −80°C until further processing. Pellets were then resuspended in 100 µL of 50% trifluoroethanol (TFE) in PBS and subsequently lysed by five freeze-thaw cycles. Lysis was further ensured by four ten-second cycles of probe-less sonication at maximum intensity (VialTeeter, Hielscher Ultrasound Technology). Proteins were solubilized by incubating the samples at 60°C for two hours. The proteins were denatured with the addition of dithiothreitol (DTT) to a concentration of 5mM and incubated for thirty minutes at 60°C, and then alkylated with the addition of iodoacetamide (IAA) to a concentration of 25 mM and incubated for another thirty minutes at room temperature in the dark. The proteins were diluted 1:5 with 100 mM ammonium bicarbonate and digested overnight at 37°C following the addition of 1 µg of Trypsin/Lys-C protease mixture (Promega) and calcium chloride to a concentration of 1 mM. The following morning, formic acid was added to each sample to a concentration of 1% and the undigested tissue was pelleted for 10 minutes in a centrifuge at 4°C. The supernatant was lyophilized and the resultant dried peptides were resuspended in 0.1% trifluoroacetic acid. The peptides were purified and desalted C18-based solid-phase capture and the eluted with 80% acetonitrile, 0.1% trifluoroacetic acid in water. Eluted peptides were once again lyophilized and resuspended in 25 µL of 0.1% formic acid. Peptide concentrations were determined by Nanodrop Lite spectrophotometer.

Two micrograms of peptide were loaded onto a 2 cm trap column (Thermo Scientific) using an Easy1000 nanoLC (Thermo Scientific). The peptides were separated and detected along a four-hour reversed-phase gradient using a 50cm EasySpray analytical C18 column coupled by electrospray ionization to a Q-Exactive HF Orbitrap mass spectrometer (Thermo Scientific) operating in a Top 25 data-dependent acquisition mode. MS^1^ data was acquired at a resolution of 120,000 with an AGC target of 3e6 ions and a maximum fill time of 240 ms. MS^2^ data was acquired at a resolution of 30,000 with an AGC target of 2e5 ions and a maximum fill time of 55 ms. A dynamic exclusion of 60 seconds was enabled, and the normalized collision energy was set to 27%.

The acquired raw data was searched using Maxquant (version 1.5.8.3) against a UniProt complete human protein sequence database (v2018_10). Two missed cleavages were permitted along with the fixed carbamidomethyl modification of cysteines, the variable oxidation of methionine and variable acetylation of the protein N-terminus. Relative label-free protein quantitation was calculated using MS^1^-level peak integration along with the matching-between-runs feature enabling a 2 min retention time matching window. False discovery rate (FDR) was set to 1% for peptide spectral matches and protein identification using a target-decoy strategy. The protein groups file was filtered for proteins identified by a minimum of two peptides and then used to carry out further analysis. The distribution of LFQ intensities were adjusted to match that of iBAQ intensities so that LFQ values could be imputed in place of missing iBAQ intensities (Wojtowicz et al., 2016). Furthermore, quantitative comprehensiveness was increased by imputing missing quantitative values with random values from a low-end normal distribution. A total of 5,708 protein groups were quantified across 33 tumor-TME paired samples. The tumour and stroma samples belonging to patient #28 were removed from differential expression analysis since this patient was later re-classified as a case with unknown primary site.

### Tissue RNA sequencing (RNAseq)

From LCM tissue, RNA was isolated using the PicoPure RNA Isolation Kit (Thermo Fisher). RNA was treated with the RNase-free DNase Set (Qiagen) and quantified using the Qubit dsRNA High Sensitivity kit (Invitrogen). Quality was measured using both the RNA Screen Tape Assay (Agilent) in combination with the 2200 TapeStation Nucleic Acid System. RNA-sequencing (RNAseq) analysis was performed at the Ontario Institute of Cancer Research (Ontario, Canada) as described previously (Connor et al., 2019). Briefly, RNA libraries were prepared using the TruSeq RNA Access Library Sample prep kit (Illumina). Library pools were quantified on the Eco Real-Time PCR Instrument using KAPA Illumina Library Quantification Kits. All above steps were performed according to the manufacturer’s protocol. Paired-end cluster generation and sequencing of 2×126 cycles was carried out for all libraries on the Illumina HiSeq 2500 platform.

Paired-end raw Fastq files were trimmed by using the following settings: HEADCROP:3 TRAILING:10 MINLEN:25 for Trimmomatic v2.38, to remove adapter content and bad quality reads. Trimmed fastqs were mapped to the human GRCH37 genome assembly using STAR v2.6.0 (Dobin et al., 2013) using the following settings: --outFilterType BySJout -- outFilterMultimapNmax 10 --alignSJoverhangMin 8 --alignSJDBoverhangMin 1 -- outFilterMismatchNmax 999 --outFilterMismatchNoverLmax 0.05 i--alignIntronMin 20 -- alignIntronMax 1000000 --alignMatesGapMax 1000000 --outFilterMatchNmin 16 -- quantMode GeneCounts --runThreadN 12. Genes quantified in at least 75% of total samples were filtered-in.

### Single-cell RNAseq analysis of human PDAC CAFs

Samples were multiplexed for single-cell RNA-sequencing using lipid-tagged indices (MULTI-seq) (McGinnis et al., 2019a). Briefly, primary human PDAC CAFs from 10 patients were trypsinised, washed twice with 1x PBS, and counted. Next, CAF single-cell suspensions (5×10^5^ cells/CAF line) were incubated with mixture of a unique barcode oligonucleotide and anchor lipid-modified oligonucleotide at 37°C for 10 minutes, followed by addition of a co-anchor and incubated for 5 minutes on ice. All subsequent steps are performed on ice or spun at 4°C. Excess barcodes were quenched with 1mL 1% BSA in 1x PBS solution in two consecutive washing steps at 400g for 5 min before cells were suspended in a final solution of 0.4% BSA in 1xPBS. Cells from all samples were pooled at equal numbers to target 2×10^4^ cells in the final transcript library. Standard protocols for the 10x Genomic Single Cell 3’ RNA kit (v3 chemistry) were used. Sequencing was performed using the NovaSeq 6000 (Illumina) as per the standard 10x configuration.

Raw sequencing reads were aligned *using CellRanger 3*.*1*.*0* with the GRCh38-3.0.0 human genome as reference. MULTI-seq data quality-control was performed using the R packages deMULTIplex v1.0.2 (McGinnis et al., 2019a), DoubletFinder_v3 (McGinnis et al., 2019b) and Seurat (v3.2.2). Following demultiplexing of the 10 samples by their unique barcodes, homotypic doublets and unlabelled cells were removed from downstream analysis using deMULTIplex. DoubletFinder was used to identify heterotypic doublets by using the homotypic doublet rate identified by deMULTIplex as input for nExp argument. Finally, only cells exhibiting <3% mitochondrial gene expression and >200 unique genes per cell were carried forth for analysis.

Cell cycle phase was annotated using the CellCycleScoring function in Seurat, with the difference between S and G2M phase scores used as input to the vars.to.regress argument within the ScaleData function. An ordered trajectory across CAF phenotypes was inferred applying the CytoTRACE computational framework (Gulati et al., 2020) to cells with >1000 unique genes. Selected genes related to CAF-activation (*FAP, ACTA2, S100A4, IL6, IL1B, NNMT)* and cell plasticity (*NT5E, ENO1, ITGB1, CD44, LGALS1, KIT)* are presented as a smoothing spline of the moving average of gene expression across 200 cells.

### Differential gene expression analysis

For human PDAC LCM tissue transcriptomes and proteomes, differentially expressed genes (DEGs) were determined using the R/Bioconductor package limma (Ritchie et al., 2015). Genes were considered as significantly regulated with an adjusted p-value threshold at 0.05 (Benjamini-Hochberg). For single cell RNAseq profiles from human PDAC CAFs, DEGs were determined using the FindAllMarkers function in Seurat using the following settings: only.pos = TRUE, min.pct = 0.25, logfc.threshold = 0.25.

### Gene set enrichment analysis (GSEA)

For human PDAC LCM tissue transcriptomes and proteomes, Fisher’s exact test was used to retrieve enriched gene sets from the lists of regulated genes. Gene sets were obtained from the Gene Ontology collection (biological process) of MSigDB v7.0 (Subramanian et al., 2005) and the ConsensusPathDB database (Herwig et al., 2016). Significance threshold was set to an adjusted p-value below 0.05. Enrichment results were visualized either as bar graphs of adjusted p-values or in Cytoscape (v3.7.1) (Shannon et al., 2003) using the Enrichment Map App (Merico et al., 2010). For single cell RNAseq profiles from human PDAC CAFs, enrichment of HALLMARK gene sets (Liberzon et al., 2015) and previously published CAF subpopulation gene sets (ecm_myCAF, TGFB_myCAF, wound_myCAF, detox_iCAF, IL_iCAF, IFNy_iCAF (Kieffer et al., 2020); iCAF and myCAF signatures (Elyada et al., 2019)) was determined using a Fisher’s exact test on markers identified for each clusters. Significance threshold was set to adjusted p-value below 0.05.

### Random Forest Model (TME PHENOtyper)

The dataset used for building the classifier included 23 subTME RNAseq profiles from reactive and deserted subTMEs (n = 11 reactive, n = 12 deserted). Data was pre-processed by centering and scaling samples via the preProcess function from the R package caret. The entire cohort was used as training dataset. The classifier was build using a random forest classification method from the randomForest R package with Leave-One-Out-Cross-Validation (LOOCV). The features selection was performed in two steps; first, an impact score was calculated for all 23,271 genes, then 72 genes with the highest score were selected (**Figure S6A**) for building the TME PHENOtyper model. Finally, a new impact score was calculated for each of these genes. To determine the robustness of the classifier against varying stroma amounts in the sample (“pseudo-bulk test”), *in-silico* samples were generated from the 23 TME samples and their paired malignant epithelium samples. These artificial samples were created by merging X% of the tumor associated sample count and 100-X% of the stroma sample count. For each pair of samples, 101 *in-silico* samples were obtained by iteratively decreasing the stroma amount by 1% from 100% (i.e. pure TME sample) to 0% (i.e. pure tumor sample). For validation analysis, the model was used to predict subTME scores from TCGA PAAD cohort and validated against histopathological assessment of main subTME phenotypes.

### TMA construction and design

Construction of the PDAC tissue microarray (TMAs) was described previously (Connor et al., 2017). Briefly, H&E sections from research paraffin blocks were reviewed by a pathologist for tumor morphology and content. An optimal area for TMA coring was marked for each block. Tissue cores (1.2 mm) were punched manually and transferred into TMA recipient paraffin blocks. Additional cores of benign pancreatic, renal, pulmonary, and hepatic tissues were included for control and TMA orientation reference purposes. For each case, multiple tumor cores were arrayed (2-4-fold redundancy) from the paraffin blocks. In total, 596 core tissue samples were obtained from 165 PDAC patients (143 treatment-naïve, 23 neoadjuvant-treated) and assembled to a tissue microarray comprising 13 slides.

### Immunohistochemistry (IHC) and semi-automated digital image analysis

Consecutive sections from above-described TMA were reviewed by a pathologist (SF) to assign the subTME present in each core and stained with Masson’s Trichrome, and Picrosirius Red and a 23-antibody panel according to standard laboratory procedures (see **Table S5** for antibody concentrations and details). For IHC, DAB+ (3,3-diaminobenzidine tetrahydrochloride) was used as a chromogen, which produces a dark brown precipitate readily detected by light microscopy. Nuclei were counterstained with hematoxylin. All slides were digitized for subsequent quantification by image analysis. Stains were quantified using QuPath bioimage analysis software (Bankhead et al., 2017). Briefly, region of Interest (ROI) grids were placed onto each slide and unique patient identifiers were superimposed onto the grids. *Simple Tissue Detection tool* was used to select for all tissue within the ROI and exclude white space. Areas containing residual normal pancreas epithelium, nerves, large blood vessels, tissue folds and stain artifact were manually excluded. Pixel-based or cell-based detection parameters were manually set and optimized for each stain. Quantification results were quality controlled by visual inspection of the 10 cores with the highest detected staining levels and the 10 cores with the lowest detected staining levels for each slide for each stain.

To account for compartment-specific expression of the markers, tumor (CK19+) and stroma (CK19-) area was measured using the *Cytokeratin Annotation* tool to obtain tumor/total and stroma/total area ratios for each core (**Figure S3A**). These were used to normalize a subset of compartment-specific stains, including fibroblast markers (FAP, α SMA) and myeloid cell markers (CD15, CD11b, CD68, CD206). In addition, the *Cell Detection* tool was used to obtain CK19+ and CK19-cell counts for each core to facilitate estimation of stromal cellularity. For ki67 staining, a classifier was created to annotate cells as either tumor or stromal. Briefly, the *Cell Detection* method was run to select all cells in the tissue. A Random Forest Classifier was then trained based manual annotations of tumor or stroma and used to finally enumerate ki67+ tumor cells. CD3+ and CD8+ tumor infiltrating lymphocytes (TILs), defined as a CD3+ or CD8+ cell in direct contact by at least one dimension with a tumor cell, were manually quantified by a pathologist (PrB). The number of CD8+ TILs was normalized by tissue area in µm2 calculated by QuPath. For analysis of GATA6 and CK5 expression, tumor epithelium was manually annotated by a pathologist (SF). The *Cell Detection* tool followed by cell-based quantification then yielded the number of negative, and weakly, moderately, and strongly positive cells.

### Second Harmonic Generation (SHG) Microscopy

SHG images were captured from unstained, uncovered TMA slides. Cores were imaged at the Advanced Optical Microscopy Facility (AOMF, UHN, Toronto) for 2 channel SHG and autofluorescence using 20x water immersion lens. Following a 1-hour laser warm-up period, a Chameleon Discovery (Coherent) femtosecond multiphoton laser was used to emit an excitation wavelength of 840 nm onto tissue. Light was collected in the backwards direction using a narrow band pass filter of 415-424 nm, capturing the SHG signal at 420 nm, half excitation wavelength. Emission was collected using a 20x water immersion objective (W Plan-Apochromat NA = 1.0). To increase uniformity of illumination across the field of view, images were acquired at zoom 2.0, generating 512×512 pixel images that were tiled to create a final image of 5120×5120 pixels (ZEN Black, ZEISS). Autofluorescence was detected at 463-617 nm and was used to provide context to the SHG image generated. SHG images were normalized via a user-defined macro for Fiji (ImageJ). Briefly, raw SHG .czi files were imported into Fiji using Bio-Format importer. Images were split into SHG and autofluorescence channels. SHG channel images were normalized to the brightest pixel in the image using Enhance Contrast, Saturated = 0.35. Images were then smoothened using a Gaussian Blur, sigma = 1. In order to remove background pixels, a Huang dark auto-threshold was applied to select only pixels positive for SHG signal. Normalized SHG images were saved as 16-bit .TIFF files. Cores with less than 25% area were omitted from further analysis. Collagen fibres detected in normalized images were quantified for alignment, straightness, width, and length, using CT-FIRE (http://loci.wisc.edu/software/ctfire) in batch-mode with parallel processing using default settings.

### Establishment and culture of PDAC patient-derived organoids (PDOs)

This study used 5 human PDAC PDO models. PDO models were generated by the Princess Margaret Living Biobank core facility using previously described protocols (Boj et al., 2015). Briefly, fresh PDAC tissue was cut into small pieces and washed with ice cold phosphate buffered saline (PBS). Tumor tissue was further dissociated to single cells or small clumps of cells using Liberase^™^ TH (Sigma Aldrich) in Advanced DMEM (Gibco) for 1 hour followed by 10-minute incubation with TrypLE Express (Invitrogen). Dissociated cells were collected, embedded in 100% growth factor-reduced Matrigel (VWR), plated in 24-well tissue culture plates as Matrigel domes and overlaid with growth medium (Boj et al., 2015). PDOs were maintained in maintained at 37°C in 5% CO2. The identity of PDOs was authenticated by short tandem repeat (STR) analysis and matched to patient tissue. PDO cultures were tested routinely for mycoplasma contamination.

### Isolation and culture of primary human PDAC CAF cultures

Fresh human PDAC tissues from 13 patients were minced into 1 mm^3^ pieces using a scalpel. Several pieces were placed in 4% formalin for subsequent paraffin embedding and histopathological assessment to determine subTMEs of the originating tissue specimens. In parallel, 5-8 pieces were plated per well on a 6-well plate in 800 µl isolation medium (DMEM/F12, 20% FCS, GlutaMAX, HEPES, Anti/Anti). Plates were placed in an incubator and media were replaced the next day and every 48 h thereafter. After 7-22 days, outgrowth of fibroblasts was observed. Cells were passaged using 0.05% trypsin when fibrotic septae formation began. Subsequent culturing was done in FCS-reduced growth medium (DMEM/F12, 8% FCS, GlutaMAX, HEPES, Anti/Anti). Cells were passaged subconfluently in fixed intervals 2 times per week using 0.05% trypsin.

### Migration Assays

CAFs were trypsinised and washed extensively in PBS to remove residual FCS. CAFs were spun down, resuspended in serum-free growth medium (DMEM/F12, GlutaMAX, HEPES, Anti/Anti), and counted with a Haematocytometer. 30,000 viable CAFs cells were seeded in triplicates into 8 μm TransWell inserts, transferred to 24-well plates containing growth medium plus 10% FCS. Cells were left for 16 h to migrate through the TransWell membrane. Plates were then removed from the incubator and media was removed from inserts. A cotton tip was used to wipe the remaining cells and excess media from the top of the TransWell membranes. Transmigrate cells at the bottom of the membrane were fixed in −20°C MeOH for 10 minutes and left to dry. Inserts were then stained overnight in 0.2% Crystal Violet (w/v) in MeOH. Inserts were washed and dried. Images of full inserts were taken with an upright brightfield microscope and analysed using the *cell counter* plugin of the ImageJ software. Data was averaged over 3-4 biological replicates and plotted +/-standard error of means (SEM).

### CAF conditioned media experiments

CAFs were seeded at 1×10^6^ cells per 10-cm plate, washed 3x with PBS 8 h later, and 5 ml Advanced DMEM/F12 medium (Gibco) was added per plate. 48 h later, conditioned media was collected, pooled, passed through 0.22 μm syringe-top filters, spun down, snap frozen and stored at −80°C. “CAF conditioned PDO media” were prepared using this CAF-conditioned Advanced DMEM/F12 as base medium for standard PDO culture medium (Boj et al., 2015). PDOs were kept in CAF conditioned PDO media for 2 passages prior to entering growth and drug response assays as well as during the assay periods.

### In vitro drug tests

CAF CM pre-educated PDOs were dissociated into single cells, counted and plated in 10% Matrigel slurries made up in CAF conditioned PDO media. PDOs were plated at 2,000 cells per well in 384-well plates in triplicate for 72 hours prior to drug treatment. Organoids were treated with a range of Gemcitabine concentrations (0.0005 −1 μM) for 96 hours. ‘Untreated’ control cells received vehicle alone (DMSO). Cell viability was determined by Celltiter Glo 3D assay according to the manufacturer’s protocol. Drug response curves were graphed using Graphpad Prism 6.0.

### Proliferation Assays

For CAF growth assays, 2,000 viable CAFs were seeded in 100 μl growth medium on 96-well plates in pentaplicates for a reference plate and five timepoints. For PDO growth assays, 2,000 viable CM-pre-educated PDO cells were plated in CAF conditioned PDO media on 384-well plates in pentaplicates. For all assay seeding, cell viability was determined by Trypan Blue exclusion on an automated Countess Cell Counter (Invitrogen). Viable cell counts were then manually confirmed using haematocytometers. At each assay timepoint for CAFs and PDOs, pre-aliquoted Cell TitreGlo 3D reagent was freshly thawed and equilibrated to room temperature for 15 minutes in the dark. Next, 1 volume of Cell TitreGlo reagent was added to the media in each well, the plate was covered, placed on a rotation shaker for 5 minutes at 120 rpm, and left at room temperature for an additional 25 minutes. Luminescence was read on a plate reader (Tecan) over 0.1 sec. Data was normalized to the t = 12h reference plate and averaged over 3-4 biological replicates for each time point before plotting mean +/- standard error of means (SEM).

### Statistical Analysis

Unless stated otherwise, bioinformatic and statistical analyses and plotting were performed using R (v3.5.1). Data were visualized using R packages ggplot2, ggpubr, gtools, reshape2, pheatmap, complexHeatmap, RColorBrewer, scales, corrplot, viridis, psych, openxlsx, dplyr. Qualitative variables were compared by Fisher exact test, and quantitative variables by Wilcoxon rank sum test for pairwise comparisons and the Kruskal–Wallis test for multiple group comparisons. Survival curves were plotted by the Kaplan–Meier method using the Survminer R package and p-values were assessed based on logrank test. The specific statistical tests used are indicated in the figure legends. All tests were two-sided. Multiple test P values were adjusted using Benjamini and Hochberg method for independent tests or Benjamini and Yekutieli method for dependent tests, respectively. Correlation coefficients were determined by the Spearman method. Statistical significance was set at p < 0.05.

### Data Availability

Raw tissue RNAseq and whole genome sequencing data will be available from the European Genome Phenome Archive (https://www.ebi.ac.uk/ega/home) under accession ID#EGAS00001002543 upon publication. Raw single cell RNAseq data will be available from the Gene Expression Omnibus platform (https://www.ncbi.nlm.nih.gov/geo/) under accession ID# GSE166571 upon publication. Raw mass spec data was deposited in UCSD’s MASSive database under accession ID# MSV000086812 and will be available from ftp://massive.ucsd.edu/MSV000086812/ upon publication. Processed data sets, scripts and data used for building the TME PHENOtyper Random Forest model are available before publication upon request.

## Notes

### Competing Interest Statement

The authors have declared no competing interest.

