## Supplemental Figures S1-S6 for "Spatially confined sub-tumor microenvironments orchestrate pancreatic cancer pathobiology"

### Supplementary Figures

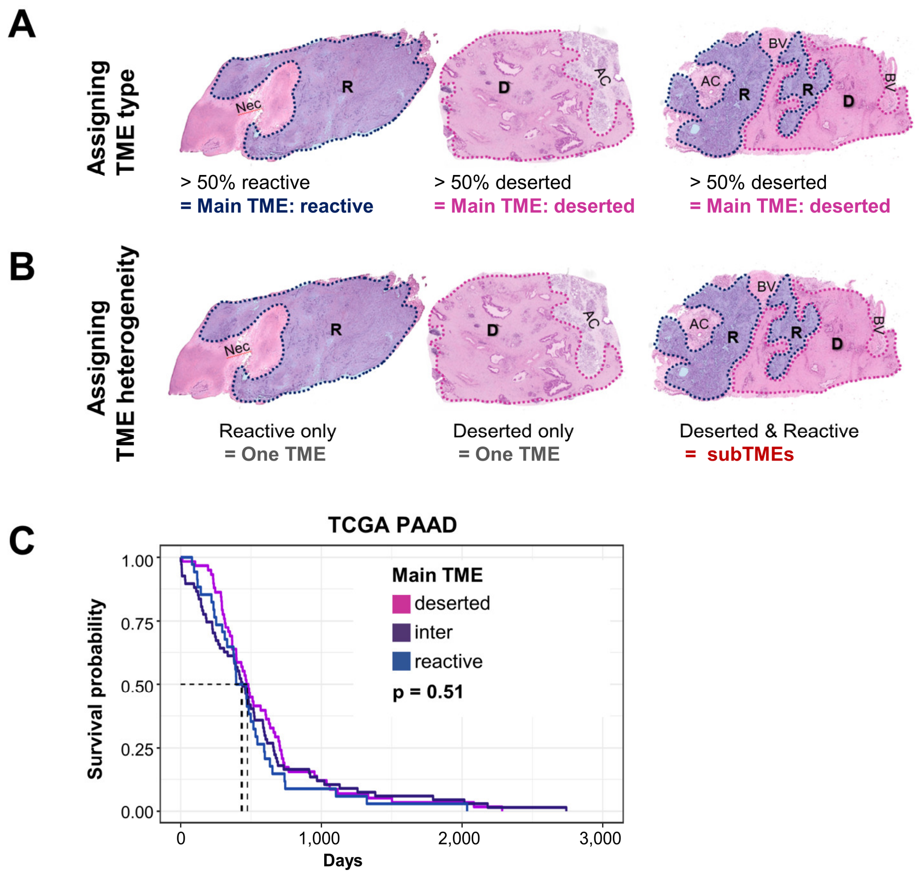


**Figure S1. Assessment of stromal phenotypic heterogeneity and predominating stromal phenotypes in human PDAC.**

Related to **Figure 1**.

**A)** Assigning main TME phenotype in PDAC patients by subTME with >50% area contribution (= main TME).

**B)** Assigning TME heterogeneity in PDAC patients as either a single dominating subTME (= One TME) or co-occurrence of regional TME phenotypes (= subTMEs).

**C)** Kaplan-Meier analysis of overall survival. TCGA PAAD cohort; strata: main TME phenotype; Log rank test.

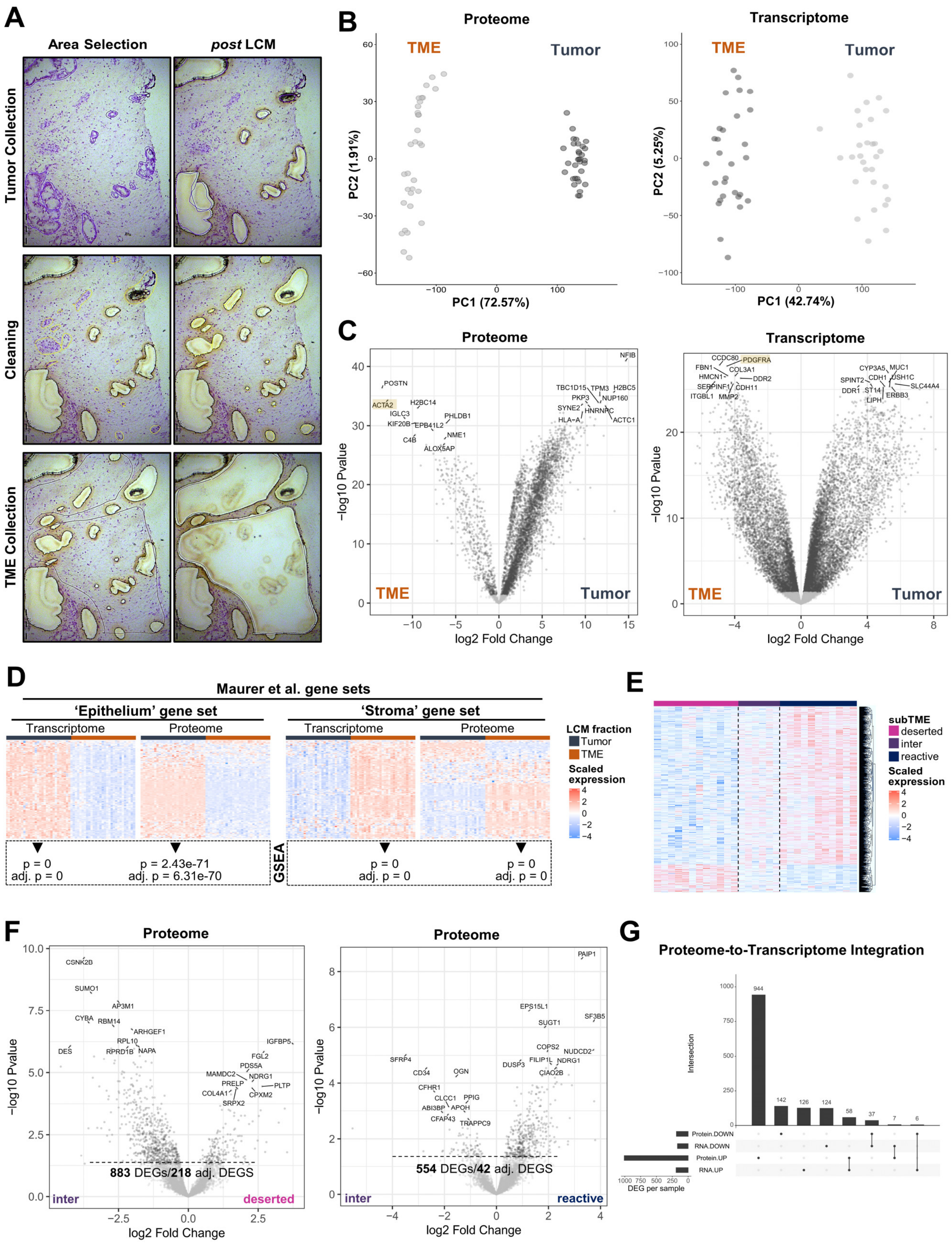


**Figure S2** – *legend on next page.*

**Figure S2. Molecular profiling of subTMEs and patient-matched malignant epithelium in PDAC.**

Related to **Figure 2**.

**A)** Laser Capture Microdissection (LCM) process for collection of PDAC tumor and TME. Representative images. Collection of malignant epithelium (*top*), followed by cleaning step to dissect and discard any non-TME elements (residual normal, large blood vessels, nerves, etc.; *middle*), and collection of subTME-specific TME fractions (*bottom*).

**B)** Principal Component Analysis (PCA) comparing human PDAC TME and tumor epithelium on the protein (*left*) and on RNA (*right*) levels.

**C)** Volcano plots: differentially expressed genes (DEGs) between human PDAC tumor epithelium and TME on the protein (*left*) and on RNA (*right*) levels.

**D)** Gene Set Enrichment Analysis (GSEA) of Epithelium and Stroma gene sets from (Maurer et al., 2019). Heatmap showing expression of gene sets (*top*). GSEA p values and adj. p values of stroma gene sets in LCM-TME profiles and epithelium gene set in LCM-tumor profiles, respectively (*bottom, in dashed outline*).

**E)** Heatmap showing DEGs (adj. $p < 0.05$) in reactive compared to deserted subTME proteomes.

**F)** Volcano plots of differentially expressed genes (DEGs) for intermediate *vs.* deserted subTMEs (*left*) and intermediate *vs.* reactive subTMEs (*right*).

**G)** Upset plot of DEGs (proteome versus transcriptome level) between stroma extracted from deserted and reactive TMEs in human PDAC.

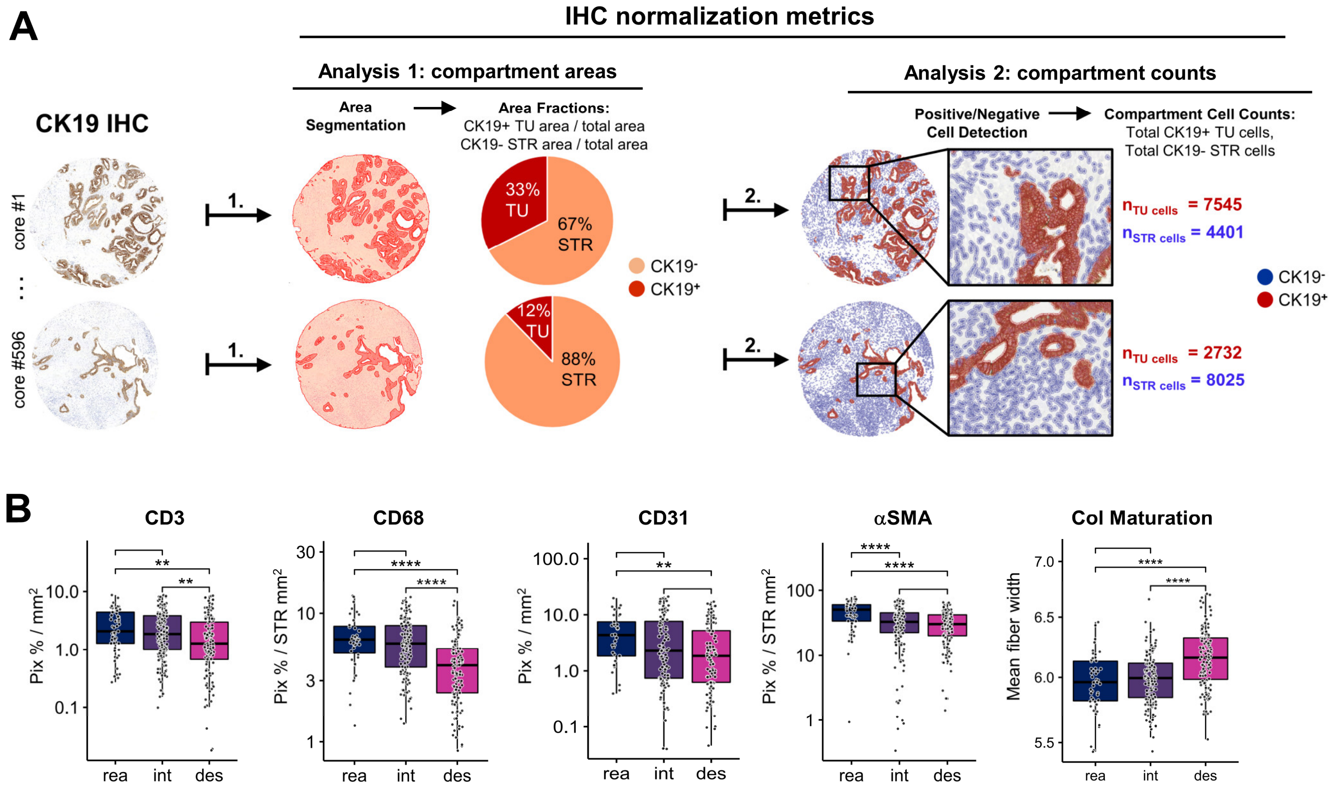


**Figure S3. Tissue profiling of human PDAC subTMEs**

Related to **Figure 3**.

**A)** Experimental schematic for assessment of tumor (TU) and stromal (STR) area (*left*) and of TU and STR cell counts (*right*) across all 596 TMA cores. Specimens were stained by CK19 IHC and digital image analysis was applied to **1.** segment images into CK19+ versus CK19- regions (*left*) and **2.** annotate cells as CK19+ versus CK19- (*right*).

**B)** Quantification of stromal composition markers CD3, CD68, CD31, αSMA stained by IHC and digitally quantified. Collagen maturation as per fibre width assessed through second harmonic generation microscopy. Boxplots compare reactive (rea), intermediate (int), and deserted (des) subTME regions. Wilcoxon test. $^{**}p < 0.01$, $^{****}p < 0.0001$

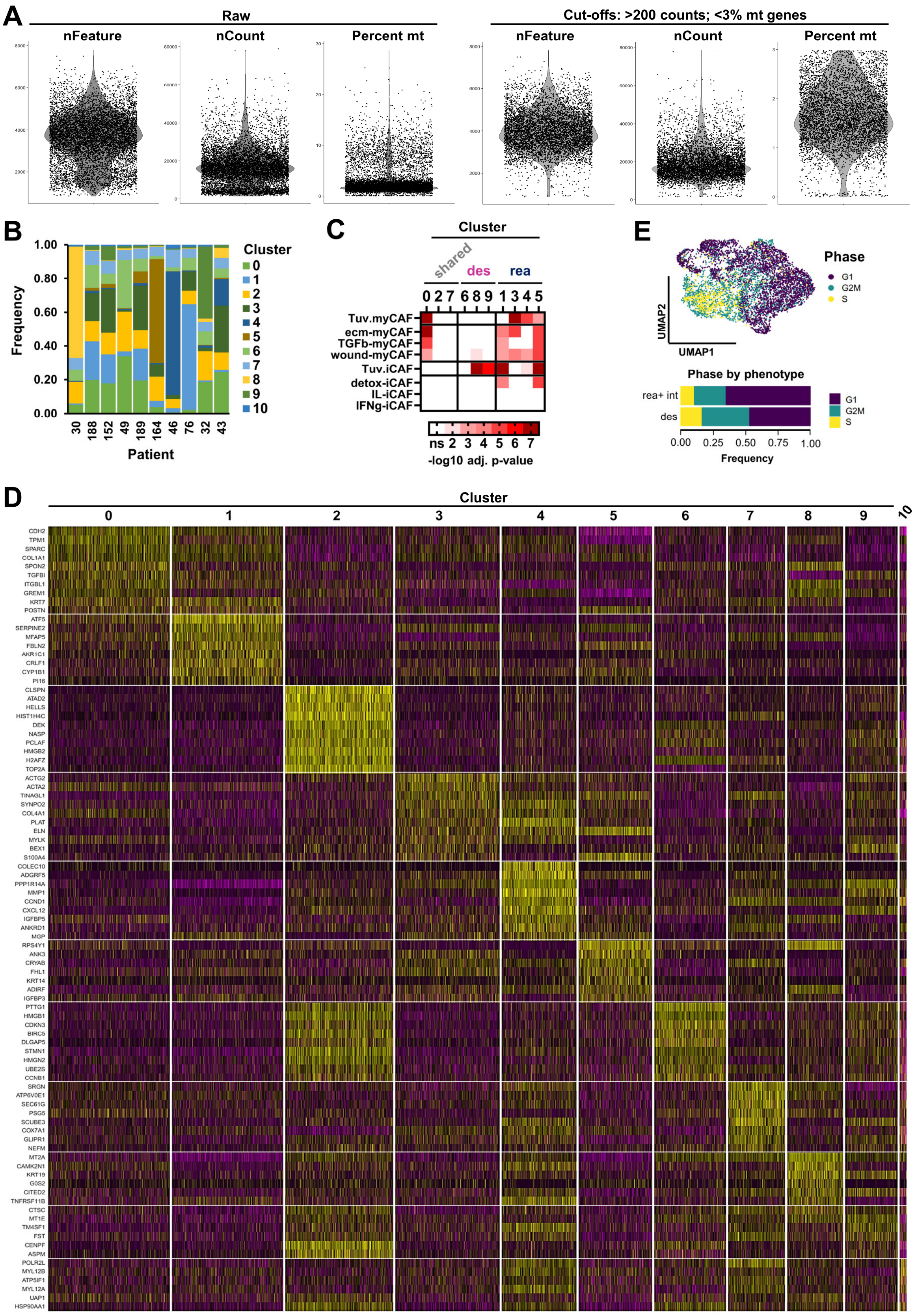


**Figure S4** – *legend on next page.*

**Figure S4. Single cell sequencing analysis of human PDAC subTME derived CAF models.** Related to **Figure 4**.

**A)** Quality control metrics for single cell RNAseq (scRNAseq) analysis of human PDAC subTME CAFs.

**B)** Relative frequency of scRNAseq clusters detected in human PDAC subTME CAFs per patient.

**C)** Enrichment analysis of pan-iCAF and pan-myCAF signatures from Tuveson (Tuv) laboratory (Elyada et al., 2019) and of ecm-myCAF, TGFb-myCAF, wound-myCAF, detox-iCAF, IL-iCAF, and IFNg-iCAF signatures from (Kieffer et al., 2020) in scRNAseq profiles of human PDAC subTME CAFs.

**D)** Top differentially expressed genes (DEGs) in scRNAseq clusters of human PDAC subTME CAFs.

**E)** Cell cycle analysis of scRNAseq profiles of human PDAC subTME CAFs. Assignment of phase in UMPA visualization (*top*). Relative frequency of phases per originating subTME phenotypes reactive (rea), intermediate (int), and deserted (des) (*bottom*).

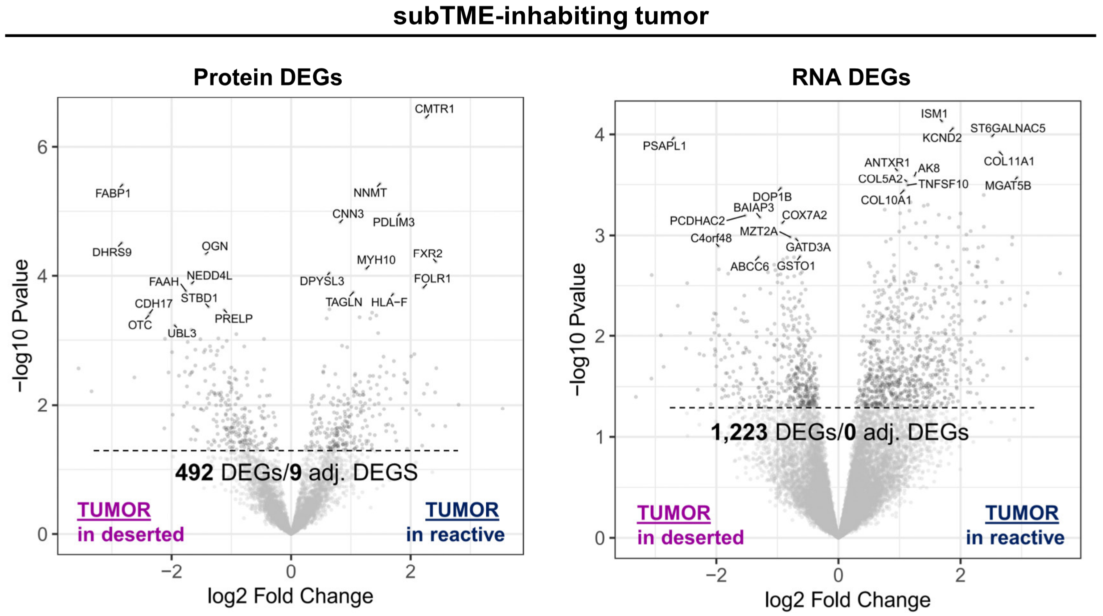


**Figure S5. MultiOMIC profiling of malignant epithelium extracted from different human PDAC subTMEs.**

Related to **Figure 5**.

Volcano plots of differentially expressed genes (DEGs) between malignant epithelium extracted from deserted versus reactive subTME regions in human PDAC. Proteomes (*left*) and transcriptomes (*right*).

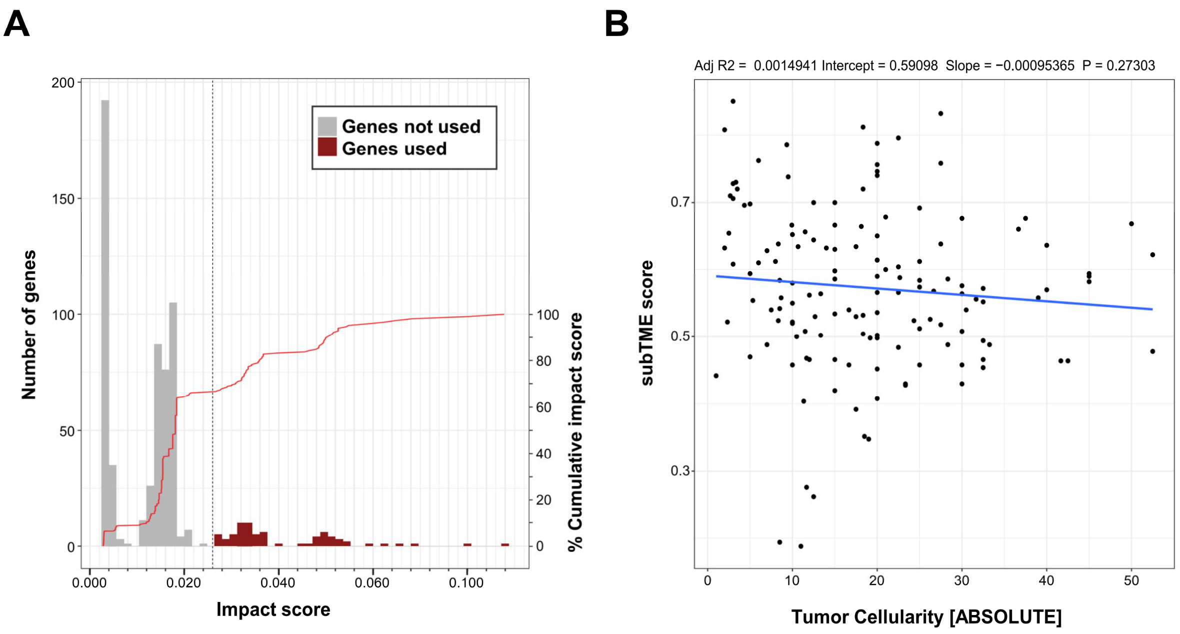


**Figure S6. Training of a Random Forest model for prediction of subTMEs from tumor bulk RNAseq profiles.**

Related to **Figure 7**.

**A)** Selection of features (n = 72) by impact score for TME PHENOtyper.

**B)** Correlation plot of TME PHENOtyper subTME scores and ABSOLUTE-based (Carter et al., 2012) estimates of tumor cellularity in TCGA PAAD cohort (n = 177)
